# Combinatorial transcription factor binding encodes cis-regulatory wiring of forebrain GABAergic neurogenesis

**DOI:** 10.1101/2023.06.28.546894

**Authors:** Rinaldo Catta-Preta, Susan Lindtner, Athena Ypsilanti, James Price, Armen Abnousi, Linda Su-Feher, Yurong Wang, Ivan Juric, Ian R. Jones, Jennifer A. Akiyama, Ming Hu, Yin Shen, Axel Visel, Len A. Pennacchio, Diane Dickel, John L R Rubenstein, Alex S Nord

**Affiliations:** Department of Neurobiology, Physiology and Behavior, and Department of Psychiatry and Behavioral Sciences, University of California, Davis, Davis, CA 95618, USA; Current Address: Department of Genetics, Blavatnik Institute, Harvard Medical School, Boston, MA 02115, USA; Nina Ireland Laboratory of Developmental Neurobiology, Department of Psychiatry and Behavioral Sciences, UCSF Weill Institute for Neurosciences, University of California, San Francisco, San Francisco, CA 94143, USA; Department of Quantitative Health Sciences, Lerner Research Institute, Cleveland Clinic Foundation, Cleveland, OH 44106, USA; NovaSignal, Los Angeles, CA 90064, USA; Environmental Genomics and Systems Biology Division, Lawrence Berkeley National Laboratory, Berkeley, CA 94720, USA; U.S. Department of Energy Joint Genome Institute, Walnut Creek, CA 94598, USA; Comparative Biochemistry Program, University of California, Berkeley, Berkeley, CA 94720, USA; School of Natural Sciences, University of California, Merced, Merced, CA 95343, USA; Institute for Human Genetics, Department of Neurology, University of California, San Francisco, San Francisco, CA 94143, USA; Department of Neurology, University of California, San Francisco, CA 94143, USA; Octant, Inc, Emeryville, CA 94608, USA

## Abstract

Transcription factors (TFs) bind combinatorially to genomic cis-regulatory elements (cREs), orchestrating transcription programs. While studies of chromatin state and chromosomal interactions have revealed dynamic neurodevelopmental cRE landscapes, parallel understanding of the underlying TF binding lags. To elucidate the combinatorial TF-cRE interactions driving mouse basal ganglia development, we integrated ChIP-seq for twelve TFs, H3K4me3-associated enhancer-promoter interactions, chromatin and transcriptional state, and transgenic enhancer assays. We identified TF-cREs modules with distinct chromatin features and enhancer activity that have complementary roles driving GABAergic neurogenesis and suppressing other developmental fates. While the majority of distal cREs were bound by one or two TFs, a small proportion were extensively bound, and these enhancers also exhibited exceptional evolutionary conservation, motif density, and complex chromosomal interactions. Our results provide new insights into how modules of combinatorial TF-cRE interactions activate and repress developmental expression programs and demonstrate the value of TF binding data in modeling gene regulatory wiring.

## INTRODUCTION

Neurogenesis in the subpallial embryonic basal ganglia (BG) produces the cells that differentiate into GABAergic and cholinergic neurons that make up mature BG structures, as well as GABAergic interneuron populations that migrate to areas such as the cortex and amygdala^1^. Comparative analysis of BG cell types and structures suggest strong evolutionary conservation of development and anatomy across the >560 million years of vertebrate phylogenic divergence^2, 3^. Many transcription factors (TFs) have been identified that control patterning and cell type specification in the BG and the brain overall, with homeobox TF genes playing central and deeply evolutionarily conserved roles^4–7^. Homeobox TFs are a large family of proteins that include a homeobox domain capable of recognizing a target DNA motif^8^. Homeobox TFs, alongside other TF classes, bind in a combinatorial and competitive manner at cis-regulatory elements (cREs) to direct dynamic expression patterns necessary for brain development and function^9–11^. Studies in simpler organisms have revealed combinatorial homeobox TF expression codes determining neuronal identity^12, 13^. Genetic studies in mice have shown a number of homeobox TFs to be critical for all stages of mammalian Central Nervous System (CNS), including BG development^6, 14^. These studies show that many individual TFs directly activate or repress transcription in developing brain, presumably via combinatorial and context-dependent TF interactions.

While single TFs have been studied at various stages of neurodevelopment, it remains largely unknown how TFs within and across diverse homeobox and other TF families work together to establish gene regulatory interactions *in vivo* in developing mammalian brain^15, 16^. More specifically, it is unknown in the developing BG and brain overall how TFs with similar or diverse binding motifs overlap in genomic targets and which combinations of TFs bind which cREs, and if cREs that are bound by the same set of TFs have similar regulatory function and evolutionary history. Characterization of TF-cRE regulatory interactions at scale is also needed to understand how sets of TFs interact to control chromatin landscapes underlying neurodevelopment. More broadly, most combinatorial TF binding studies that compare more than a few TFs in a single system have been done in less complex organisms or *in vitro* cell models, thus there remain major questions regarding how sets of TFs bind to regulatory DNA targets during mammalian embryonic development and what the relationships are between cis-trans interactions between TFs and regulatory targets and chromosomal DNA interactions. We addressed these questions via integrating ChIP-seq data from 12 TFs, 9 of them representing 6 distinct homeobox classes, with chromatin state and chromosomal interactions identified using H3K4me3 PLAC-seq, and established the activity of representative TF-bound cREs in transgenic mouse enhancer assays (Figure 1A).

**Figure 1.**
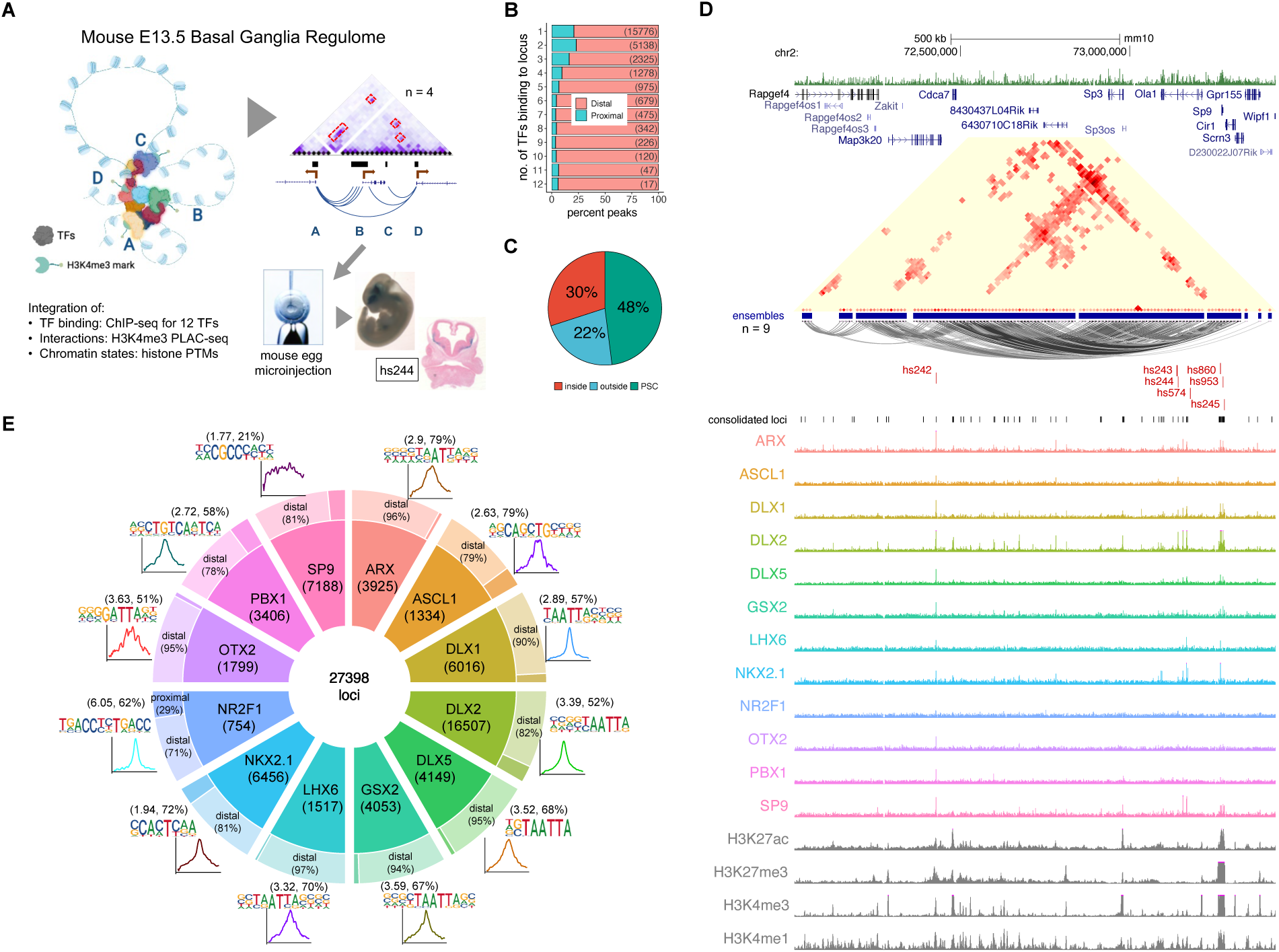
TF Binding Profiles and Basic Genomic Features. **(A)** Schematics of the scope of the present study, showing 3D structures combinatorially bound by TFs in transcriptionally active chromatin (H3K4me3-marked). The model was validated by enhancer transgenic mouse assays. **(B)** Distribution of combined bound loci between distal and proximal regions, segmented by the number of TFs sharing locus position. **(C)** Pie chart showing the distribution of loci in relation to the loops formed by H3K4me3-mediated PLAC-seq contacts (PSC). **(D)** *Sp9* locus showing TF binding and PLAC-seq interactions with VISTA enhancers *hs242*, *hs243*, *hs244*, *hs245*, *hs574*, *hs860*, and *hs953*. PLAC-seq contacts are displayed as arcs and contact maps (adj. p < 0.01). **(E)** Doughnut plot showing individual TF number of binding loci, split into proximal and distal from gene TSS, with the associated core motifs and their average distribution around peak centers. In parentheses are the enrichments over background and percent of target motifs. See also Figure S1.

## RESULTS

### A TF-anchored model of regulatory interactions in E13.5 mouse basal ganglia

We performed epigenetic experiments on micro-dissected embryonic day (E)13.5 mouse BG, when GABAergic and cholinergic neurogenesis is ongoing in the medial, lateral, and caudal ganglionic eminences (MGE, LGE, and CGE)^17^. Dissections included the ventricular zone (VZ), where neural precursor stem cells are located and expanding, the subventricular zone (SVZ), which includes intermediate progenitors and early born neurons, and the mantle zone (MZ), which is made up of maturing and migrating immature GABAergic and cholinergic neurons^17^. We performed ChIP-seq targeting five TFs (ARX, ASCL1, GSX2, NR2F1, and PBX1/2/3) and combined this new data with seven previously published BG ChIP-seq TF datasets (DLX1, DLX2, DLX5, LHX6, NKX2.1, OTX2, and SP9)^18–21^ (See Figure S1 and Online Methods for computational and experimental details). The PBX antibody used in ChIP-seq experiments detected PBX1, PBX2, and PBX3 proteins, but results here are referred to as PBX1 for simplicity. This representative TF set in this study includes: 1) TFs that establish regional identity and control proliferation and are expressed most highly in neural progenitors (ASCL1, GSX2, NR2F1, OTX2); 2) TFs that activate neurogenic transcriptional programs and are expressed most highly in the VZ-SVZ transition (DLX1, DLX2, NKX2.1), and 3) TFs that drive maturation of GABAergic neurons (ARX, DLX5, LHX6, PBX1, SP9) expressed most strongly in the SVZ and MZ, as shown by expression in single cell RNA-seq and *in situ* data (Figure S1b, S1c)^22^. In addition to diverse functions and expression patterns, these TFs capture a diverse set of TF families, including 9 homeobox TFs from 6 classes: Distal-less/DLX (DLX1, DLX2, DLX5), LIM (LHX6), HOXL (GSX2), PRD (ARX, OTX2), and NKL (NKX2.1), as well as three non-homeobox TFs, ASCL1 (bHLH family), NR2F1 (COUP orphan nuclear receptor family), and SP9 (SP family, buttonhead-like ZF).

We first identified the genome-wide targets from ChIP-seq data for each TF, and then generated a merged set totaling 27,398 loci, with each locus targeted by at least one TF (see Online Methods). The full set of merged loci with TF binding included 22,297 distal putative cREs (pREs) and 5,101 promoter-proximal sites that overlapped or were less than 2kb from a TSS. TF-bound loci were enriched near genes associated with neurodevelopment and with specific functions that spanned proliferation, neurogenesis, and neuronal maturation. Individual TF ChIP-seq peak sets varied in number of peaks, percent distal versus proximal targets, and primary binding motifs (Figure 1E). All TFs except SP9 had strong enrichment of a primary binding motif centered within ChIP-seq peaks, indicating mostly direct DNA binding. SP9-bound pREs includes a subset that contain a putative SP9 primary binding motif^23^, suggesting that while direct binding occurs, the majority of SP9 interactions here are indirect. Illustrating the challenges of inferring TF interactions via motif analysis alone, six of the TFs (ARX, DLX1, DLX2, DLX5, GSX2, and LHX6) recognize variations of a highly similar “TAATTA” motif common to many homeobox TFs^24^. Despite recognizing the same motif, genomic targets of these 6 TFs varied substantially (Figure S1e, S1f). Considering the set of loci targeted by these 12 representative TFs, the majority featured a peak call from only one TF (57.5%), though many pREs featured multiple TF peaks and a small subset (2.7%) included a ChIP-seq peak from 8-12 of the TFs (Figure 1B).

To understand the cis-regulatory contexts associated with TF binding, we integrated biophysical interactions, chromatin state, and gene expression data. We performed H3K4me3-anchored Proximity Ligation-Assisted Chromatin Immunoprecipitation followed by sequencing (PLAC-seq) at 10-kb resolution. We identified 113,048 significant interactions that represented 13,128 PLAC-seq contacts (PSCs). These interactions comprise 1,974 distinct “ensembles” built from chaining together inclusive sets of interacting regions (see Online Methods). Most of the interaction ensembles included only one or two PSC interactions, however, there were a substantial number of complex ensembles with 5 or more chained PSCs. Many of these complex ensembles featured extensive interactions between distal and proximal contacts, and nearly all had overlap with at least one TF-bound pRE (Figure S1g, S1h). Across TF-bound pREs, 48% overlapped PSCs and 30% were within loops formed by these contacts (Figure 1C). Via PLAC-seq interactions, we were able to map nearly half of the TF-bound pREs to putative regulatory target genes and evaluate relationships between TF-binding and interaction structure. We additionally segmented the genome into 9 chromatin states via ChromHMM using ChIP-seq data for H3K4me3, H3K4me1, H3K27ac, and H3K27me3 (Figure S1i, S1j). Lastly, we associated published E13.5 BG RNA-seq to complete the landscape. As expected, PSCs were biased towards chromatin states featuring H3K4me3 and were associated with increased gene expression, but were also enriched for states with H3K27ac, H3K4me1, and H3K27me3, indicating chromosomal interaction data captures transcriptionally active and bivalent enhancer-promoter complexes. The *Sp9* locus is an example showing the intersection of TF binding, chromosomal interactions, and chromatin state (Figure 1D). Together, these datasets synthesize regulatory interactions between cREs and TFs, offering an integrated map of the regulome in E13.5 mouse BG.

### Combinatorial TF binding defines distinct enhancer and promoter pRE sets

We next sought to define combinatorial binding patterns of the 12 TFs. Comparing peak overlap has limitations for modeling combinatorial binding due to differences in antibody and ChIP-seq performance, as well as missing differences in strength and spread of signal, and in complex binding patterns. Thus, to better model TF binding patterns, we leveraged unbiased K-means clustering implemented in DeepTools^25^, using as input normalized ChIP-seq neighborhood signal coverage across a 1-kb region centered on TF-bound loci (Figure 2A). This approach yielded 7 clusters of promoter-proximal pREs and 18 clusters for distal pREs (Figure 2B, 2C). All TFs exhibited strong enrichment in specific pRE cluster sets, accompanied by weaker or no binding in other clusters.

**Figure 2.**
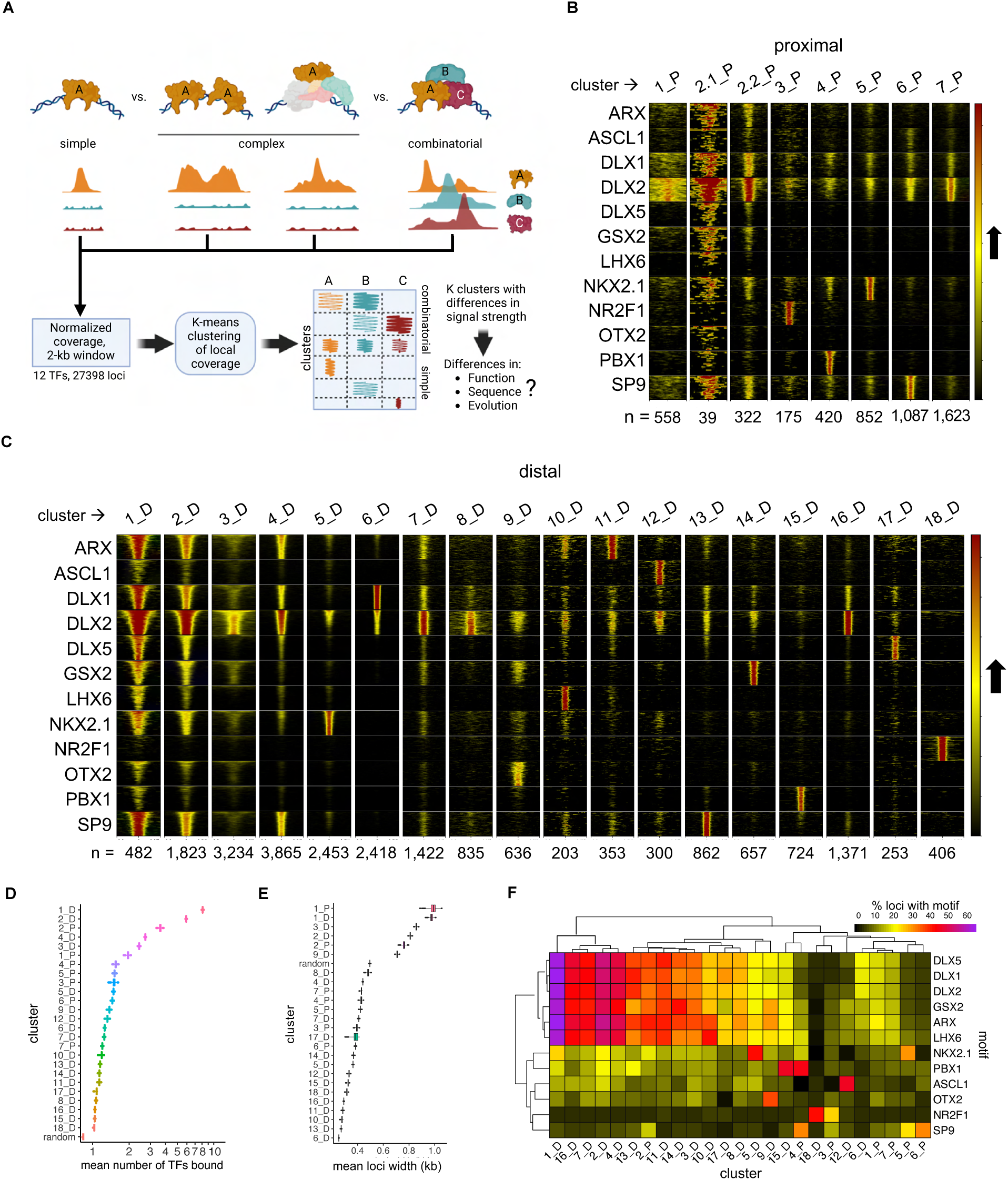
Organization of Bound Loci into Clusters of Similar Binding Neighborhood Profiles. **(A)** Schematics showing the several binding patterns captured by investigating local neighborhood around ChIP-seq peak summits, as well the subsequent clustering and following genomic profile characterization. **(B)** and **(C)** Heatmaps representing each TF coverage around 1 kb of each called peak in proximal and distal regions, respectively. Row blocks and columns depict TFs and clusters, respectively. Within each row block, each line represents the coverage color-codes for intensity of ChIP-seq signal (intensity grows in the black-to-yellow-to-red direction). Within each cluster, lines with same position across the TFs represent the same genomic locus. Numbers (n =) indicate the number of peaks called within each cluster. **(D)** Distribution of mean number of TFs bound to each locus across clusters. Mean distributions were calculated by sampling without replacement (N=1000). Random means was calculated by randomly sampling the genome (N=27398). **(E)** Distribution of mean number of TFs sharing loci across binding clusters, and compared to a random sample, calculated by sampling without replacement (N=1000). **(F)** Heatmap showing the relative enrichment of core binding motifs for each of the TFs across clusters. See also Figure S2.

There were two general binding patterns for pREs: clusters with one or two dominant TFs and an average of 1 to 2 ChIP-seq TF peaks versus clusters with multiple TF binding (“broadly-bound”) and an average of greater than two TF ChIP-seq peaks. Clusters 1_D and 2_D had the highest TF co-occupancy, with an average of 6-8 of the 12 TFs bound per pRE (Figure 2D): DLX2 was the only TF where ChIP-seq signal was widespread across most clusters, though binding intensity varied and was absent in some clusters (e.g., 15_D and 18_D, which were exclusively bound by PBX1 and NR2F1, respectively). NR2F1 was the only TF whose proximal and distal interaction sets (3_P and 18_D) did not have a clear overlap with other TFs. Average pRE width differed by cluster (Figure 2E), which was driven by increased ChIP-seq signal intensity and/or local spread. Cis-level motif occurrences in DNA across pRE clusters mirrored TF binding (Figure 2F). For example, there was high specificity between the ChIP-seq binding for relevant TF and presence of DNA motif for ASCL1, NKX2.1, NR2F1, PBX1, and OTX2. There was expected high correlation across TAATTA motif variants for the set of TFs that bind these motifs (ARX, DLX1, DLX2, DLX3, GSX2, and LHX6), with highest rate of TAATTA motif occurrence in broadly-bound clusters 1_D and 2_D. Cluster 6_D featured DLX1 binding and was strongly enriched for tandem TTAA simple repeats, consistent with repeat element binding or the “decoy” model of gene expression regulation^26^. We additionally tested all HOMER motifs and found numerous motifs with cluster-specific enrichment, for example Foxo1, Oct4, and Sox2 motifs in 1_D and 2_D (Figure S3b)^27^.

In summary, clustering by local ChIP-seq neighborhood signal separated TF targets into distinct pRE groups with specific combinatorial TF binding signatures. pRE sequences were generally enriched for the cognate motifs of bound TFs, with the expected exception of SP9. While we expected to identify various discrete combinatorial binding patterns, somewhat surprisingly, 8% of distal pREs that comprised clusters 1_D and 2_D were broadly bound across TFs rather than specific to a particular TF subset. Most strikingly, the small set of distal loci making up 1_D were bound by nearly all TFs, with representation across all 9 homeobox TFs, and were further distinguished by representing the extremes for ChIP-seq signal for TAATTA-binding homeobox TFs.

### Combinatorial TF binding reveals cREs with distinct neurodevelopmental roles, cell-type specificity, and regulatory function

We next tested if pREs with different TF binding patterns similarly exhibit differences in chromatin state, regulatory targets, cell-type specific chromatin accessibility, chromosomal interactions, and sequence features (Figure 3). We compared chromatin states across pRE clusters (Figure 3A). Proximal pRE clusters were enriched for active and bivalent promoter states (H3K4me3 with or without H3K27me3), consistent with constitutive and developmentally-regulated promoters. Distal pREs sets separated into groups with differential representation of enhancer-relevant states: no histone marks or repressed (H3K27me3), active or bivalent (H3K27ac without or with H3K27me3), inactive or poised (no marks or H3K4me1 without H3K27ac), and a mix of active, poised, and repressed states. Clusters featuring a single TF were more likely to exhibit inactive or repressed states, whereas broadly-bound pREs were active or bivalent. Exceptions were 12_D, with ASCL1-specific TF binding and active enhancer states, and 4_D, with binding across several TFs but inactive/poised states.

**Figure 3.**
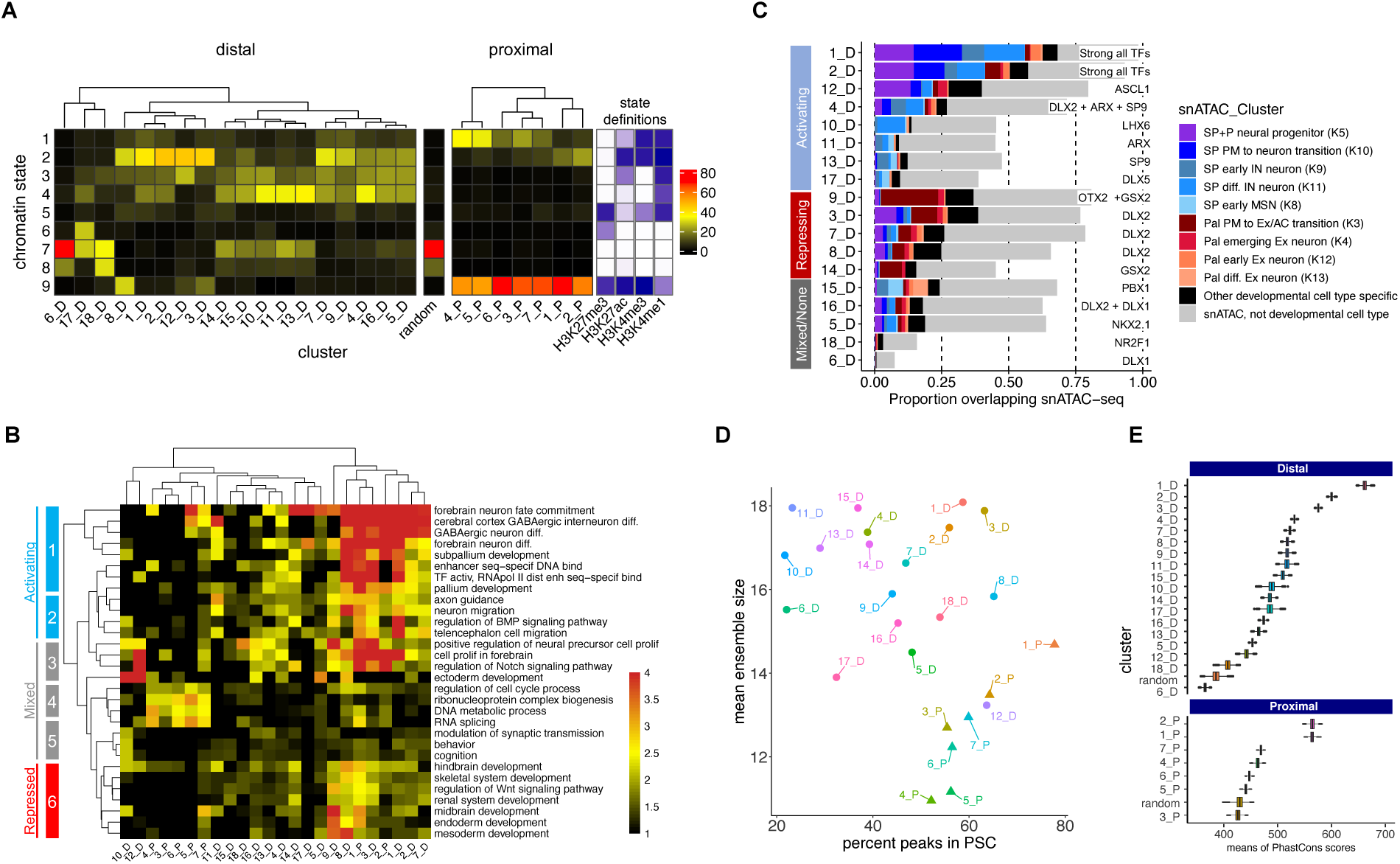
Genomic and Functional Features of TF-Bound RE Clusters. **(A)** Frequency of occurrence of peaks across chromatin states and binding clusters, split into distal and proximal. As reference, in between the two heatmaps is one derived from assigning chromatin states to a random loci sample. Color codes represent the percentage of peaks by cluster. **(B)** Gene ontology analysis of genes hitting interaction contact points (PSC) split into clusters, displaying select brain-specific or general terms. **(C)** Bar plot depicting the intersection of our binding clustering with clusters determined by single-nucleus ATAC-seq^28^, showing putative neuronal cell differentiation states across binding clusters. **(D)** Dot plot showing the mean ensemble size across binding clusters in function of the percent of loci in the cluster colocalizing with PSCs. **(E)** Distribution of means of PhastCons scores across clusters for 60 vertebrates. Top panels are distal peaks, and proximal ones are on the bottom panels. References were random distal and proximal genomic regions of random widths. See also Figure S3.

We used a neighborhood-based approach (GREAT) for target gene assignment and functional annotation enrichment analysis of pRE clusters (Figure 3B). PLAC-seq defined gene target assignment showed overall agreement (Figure S3a). Beyond shared general enrichment for neurodevelopmental pathways, contrasts emerged in target genes and pathways across TF-pRE sets. Combining chromatin state-based activity inference and pathways enrichment revealed signatures of activating, mixed activating/repressive/inactive, and repressive TF-cRE modules. Broadly-bound pREs, both distal (1_D, 2_D, 3_D, 7_D, and 8_D) and proximal (1_P and 2_P) regulated a specific program of subpallial development and GABAergic neuron differentiation. These pREs target genes regulating GABAergic neurogenesis and associated processes such as axon guidance and migration. Contrasting this activation, TF-pRE modules 9_D, 3_D, 8_D, and 1_P repress transcriptional programs associated with earlier embryogenesis, as well as other organ systems and CNS structures. For example, repression of skeletal and renal systems and endoderm and mesoderm programs. We identified mixed activation and repression of neural precursor proliferation directed by specific TF-pRE binding. For example, ASCL1-specific pREs (12_D) activated neural precursor expansion, while 9_D, 3_D and 8_D were associated with bivalent and repressive regulatory states for interneuron differentiation. TF-specific promoter pRE modules were most strongly enriched for housekeeping functions (e.g., RNA splicing ad DNA metabolism).

We intersected pRE clusters with a published set of open chromatin regions (OCRs) annotated to specific cell types in adult and developing forebrain via single nucleus (sn)ATAC-seq^28^ (Figure 3C). 63% of pREs overlap OCRs overall, with 26% of pREs overlapping a developmental cell-type specific OCR compared to only 3% for adult cell-type specific OCRs. Neurodevelopmental OCRs separated into cell-type specific sets for neural progenitors and distinct sub-pallial (SP) and pallial (P) populations of maturing excitatory and inhibitory neurons. The two TF-pRE modules with strongly active chromatin and the broadest TF binding, 1_D and 2_D were again outliers with the highest overlap with cell-type specific developmental OCRs (70% and 60%, respectively), overlapping OCRs from combined SP/P neural progenitors and three early stages of maturing SP neurons. In contrast, 12_D, 9_D, and 3_D modules had reduced but still high neurodevelopmental OCR overlap, but with different cell-type OCR classes. 12_D was enriched for neural progenitor OCRs, while 3_D and particularly 9_D were enriched in OCRs mapping to pallial early excitatory neurons. TF-pRE clusters further showed stage specific cis-regulatory programs within the BG, for example 12_D and 3_D had enrichment for progenitor OCRs while 4_D was enriched for early and differentiating SP neuron OCRs. Overall, integration of chromatin state, functional enrichment, and snATAC-seq OCRs reveals combinatorial TF binding in embryonic BG directs activation of GABAergic neurogenesis while repressing earlier and alternative developmental programs, including repressing pallial and excitatory fates.

Intersection of pREs and PLAC-seq interaction ensembles showed differences across TF-pRE clusters (Figure 3D). Broadly-bound pRE clusters featured higher than average rates of inclusion in PSCs and participated in ensembles with increased average number of PSCs. Among interaction ensembles with more PSCs, complex interactions between distal pREs as well as distal and proximal pREs, were common, consistent with a biophysical structure model of TF-cRE biomolecular condensates in the nucleus that increase recruitment efficiency of co-factors and RNA polymerase complexes^29–31^. Lastly, we compared evolutionary sequence conservation using the maximum vertebrate Phastcons element s score for each pRE (Figure 3E). Average proximal pRE cluster scores ranged from 410, equivalent to a random sample of promoter proximal intervals, to 580 for 1_P and 2_P. Distal clusters spanned from 365, similar to randomly selected distal regions, to the maximum conservation score of 662 for cluster 1_D. There was a strong relationship between number of TFs bound and evolutionary conservation for distal pREs; again, cluster 1_D was the extreme case.

### Enhancer activity in developing mouse telencephalon is predicted by combinatorial TF binding

To assess whether distal elements bound by multiple TFs are *bona fide* enhancers active in the developing BG, we identified pREs that overlapped with VISTA enhancer elements^32^ and classified as active in SP (n=60), P (n=57), SP and P (n= 70), and non-telencephalic brain (non-tel, n=75) (Figure 4E, Supplementary Table 3). As a comparison set, we included tested VISTA elements that had no reproducible enhancer activity (n=121). Distal pREs with high levels of combinatorial binding (8-12 TFs, clusters 1_D and 2_D) overlapped 48% of all VISTA enhancers with SP activity, 33% with SP and P activity, and 25% with P activity, but only 8% of those with non-telencephalic brain activity and 3% with no activity (Figure 4A, 4B). Conversely, VISTA elements without BG TF binding made up 53% of those with non-telencephalic activity and 64% with no activity (64%). Individual TF ChIP-seq peaks showed similar enrichment for VISTA enhancers with subpallial activity versus non-telencephalic enhancers (Figure S4a, S4d). These results indicate that broadly-bound distal pREs indeed function as enhancers *in vivo* in developing mouse telencephalon, with clear enrichment for subpallial activity. Strikingly, ChIP-seq of the 12 TFs here identified most BG-active enhancers in the VISTA database, as well as many enhancers active in other tissues. This suggests that developmental enhancers feature complex TF binding driving both activation, here in embryonic BG, as well suppression of activity in other cells and tissues.

**Figure 4.**
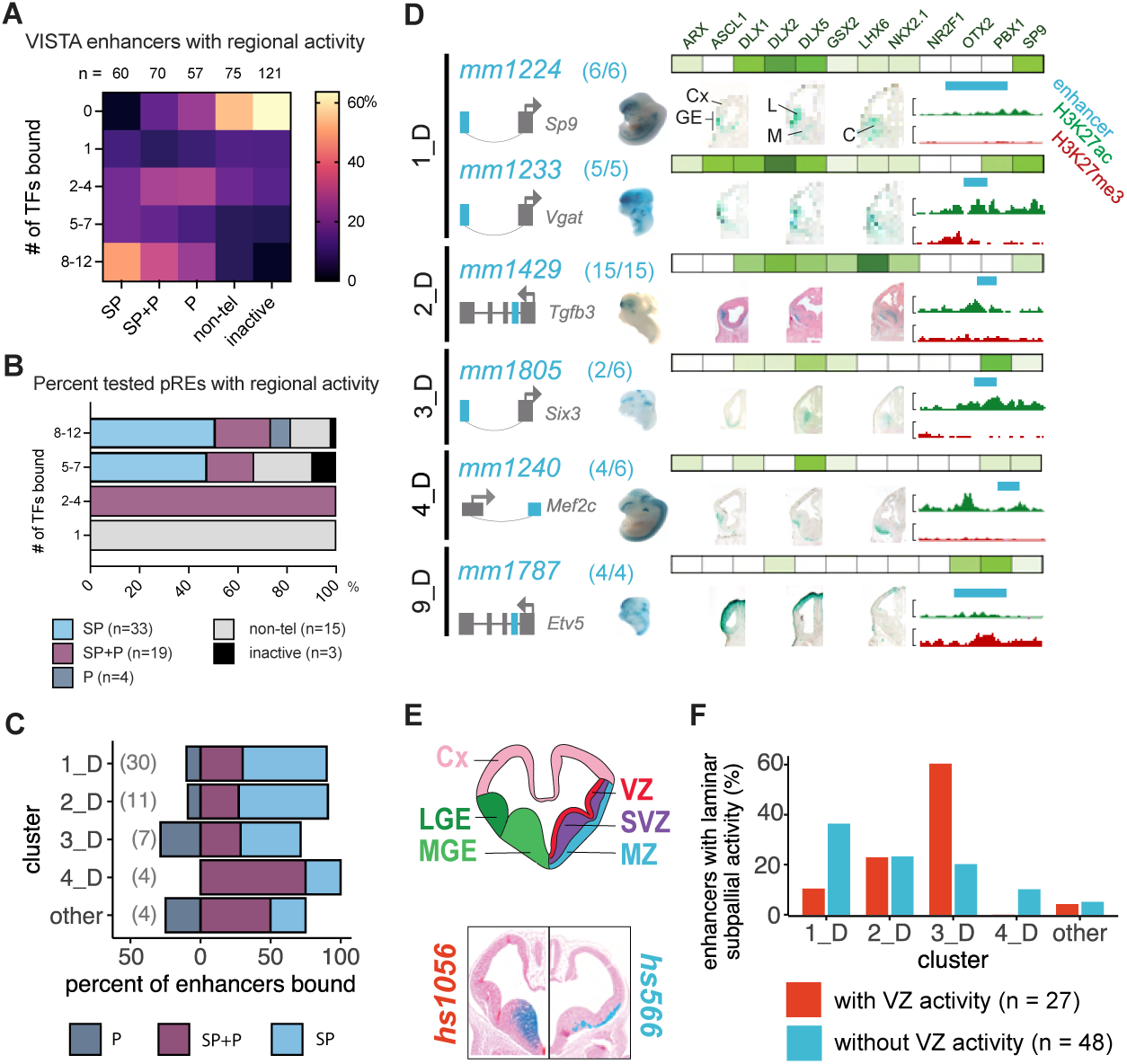
VISTA and Novel Enhancer Activity across Bound Loci. **(A)** Heatmap of combinatorial TF binding (percentage enrichment, 0-12 TFs) on VISTA enhancers that have subpallial (SP), pallial and subpallial (SP+P), pallial (P), non-telencephalic (non-tel), and no activity (inactive). **(B)** Stacked bar plot showing the percentage of newly identified regulatory sequences with high (8-12 TFs), intermediate (5-7 TFs), low (3-4 TFs), and very low (1 TF) binding in BG showing spatial regional activity. **(C)** Stacked bar plot depicting percentage of novel enhancers across binding clusters showing restricted spatial enhancer activity in the subpallium, pallium and shared among them. **(D)** Six pREs representing different clusters that were tested for activity in transgenic mouse assays^33^. Clusters are shown in the left column, enhancer names are written in turquoise, and the success ratios are listed next to the name (i.e., 6/6 depicts 6 embryos with forebrain activity out of 6 embryos tested). Schemas predict the regulated genes by the tested enhancers (turquoise). The grey arrow depicts the orientation of the TSS. Green bars show the normalized binding of BG TFs, with color intensity proportional to ChIP-seq intensity. The specific TFs bound are shown above the top bar. Wholemounts (WM) and three sections representing the LacZ expression are shown. H3K27ac (green) and H3K27me3 (red) histone ChIP-seq results from the GE are shown to the right; the turquoise bars correspond to the tested genomic regions. Cx: Cortex; GE: Ganglionic Eminences; L: LGE; M: MGE; C: CGE. **(E)** Coronal brain section schematization showing: 1. the subregions of the primordial BG (LGE and MGE) as well as the cortex (top left hemisection) ; 2. the subregional laminae of the GEs (VZ, SVZ, and MZ; top right hemisection). Hemisections from 2 VISTA enhancers with specific sub regional activity are shown below with *hs1056* showing activity in the VZ and SVZ of the MGE (bottom left hemisection) and *hs566* showing activity in the mantle zones of the MGE and LGE (bottom right hemisection). **(F)** Bar plot depicting cluster classification of enhancers with VZ and non-VZ activity (n=99). See also Figure S4.

Elements in the VISTA database were identified via criteria that may bias this set towards general enhancer activity, such as ultra-conservation^34^ and forebrain p300 ChIP-seq^35^. To directly test our TF-based pRE predictions and towards generating a resource of subpallial enhancers, we identified 84 novel TF-bound pRE loci and tested them using the same transgenic enhancer assay as used in VISTA discovery^32^. In addition to selecting candidates based on TF binding, we also chose enhancers with PLAC-seq identified target genes that play critical roles in subpallial development, enriching the value of enhancers screened here. Individual results for these enhancers are shown in Figure S4. Similar to findings for the VISTA elements, enhancer activities of newly-tested pREs with broader BG TF binding were more likely to exhibit subpallial specificity (Figure 4C). Splitting by cluster, pREs from clusters 1_D and 2_D were particularly enriched for subpallial activity, while cluster 3_D showed similar specificity in enhancer activity in the SP, SP+P and P (Figure 4D and Figure S4b, S4c). Six tested pREs representing 5 clusters are shown with ChIP-seq signal and local H3K27ac and H3K27me3 in Figure 4D. Finally, we assessed if pRE clusters had differential enhancer activity in subpallial VZ progenitors versus neurons (examples shown in Figure 4E). Cluster 1_D had proportionally higher non-VZ versus VZ enhancer activity; 2_D had a balance of VZ and non-VZ enhancer activity, and 3_D had higher VZ activity (Figure 4F), matching gene ontology analyses and snATAC-seq inferences above. In terms of individual TFs, only OTX2 and PBX1 showed preferential binding to enhancers active in the VZ (Figure S4e). ASCL1, DLX1, DLX5 and LHX6 preferentially bind to enhancers with activity in the SVZ/MZ. Thus, different TF-pREs modules exhibit differential activity in progenitor versus post-mitotic states and between telencephalon regions.

### Broadly-bound enhancers are characterized by TAATTA homeobox motif composition and deep evolutionary conservation

To test how sequence determinants distinguish the combinatorial TF binding clusters, we compared frequency of motif occurrence, spacing and position of motif pairs (i.e., motif “grammar”), and base-level evolutionary conservation (vertebrate PhyloP score) of motif and flanking DNA (Figure 5). For this analysis, we generated a merged set of TAATTA motifs. We further separated TAATTA motifs into “symmetric” instances, where the motif was identified as overlapping occurrences on sense and antisense strands (i.e., palindromic), “degenerate” instances identified on only one strand, and “complex” overlapping instances that largely map to simple “TTAA” repeats. The majority of TAATTA motifs identified in random regions are degenerate, suggesting symmetric instances are more likely to be functional. Only 6_D was enriched for complex TAATTA motifs, consistent with these pREs harboring simple TTAA repeats. The average number of TAATTA motifs within pRE had significant range across clusters, and both motif counts and relative symmetric:degenerate motif ratio showed cluster-specific patterns (Figure 5A). 1_D averaged nearly four TAATTA instances per pRE and had the largest shift towards symmetric over degenerate instances. These patterns show that 1_D represents an extreme in both binding patterns and sequence composition. The other broadly bound distal clusters of 2_D, 3_D, and 4_D also all averaged multiple TAATTA instances and increased ratio of symmetric motifs.

**Figure 5.**
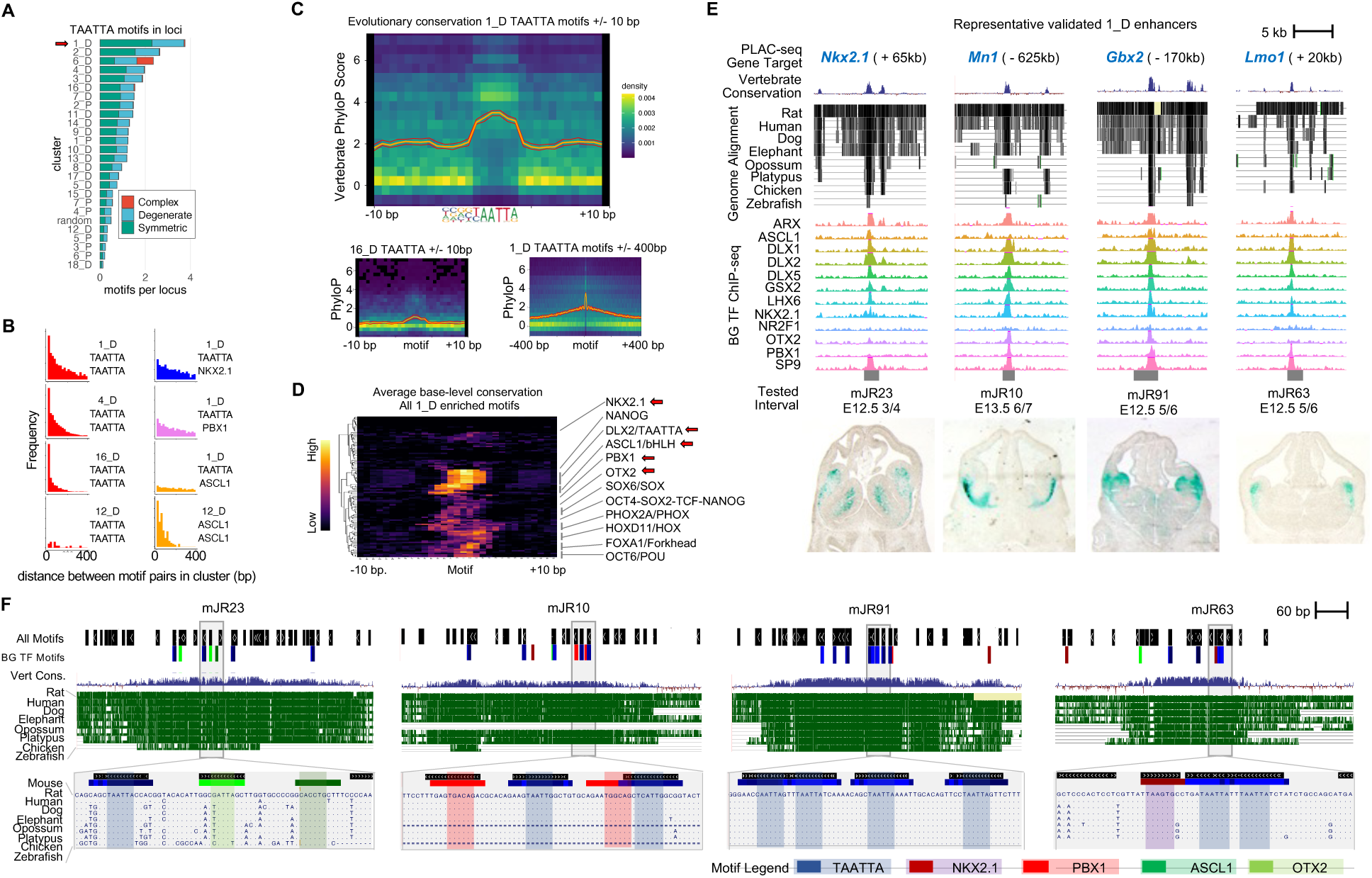
Arrayed TAATTA motifs anchor deeply-conserved GABAergic enhancers. **(A)** Relative number of TAATTA motifs within each RE across clusters separated by symmetric, degenerate, and complex instances. **(B)** Histogram showing distribution of distance in base pairs between all motif pair occurrences within REs for selected clusters and motif pairs. **(C)** Average base-level sequence conservation (vertebrate PhyloP score) for TAATTA motif and flanking DNA for 1_D (top) and 16_D (bottom left) within 10bp of motif, and for 1_D out to 400bp of motif (bottom right). **(D)** TAATTA motifs exhibit the strongest base-level conservation across TF motif families enriched in 1_D REs. Enriched motifs with significant base-level conservation increase compared to 10bp flanking sequence labeled, primary motifs from BG TFs in bold. **(E)** Four representative 1_D REs with enhancer activity. Target gene, evolutionary conservation, BG TF binding, and enhancer activity in E12.5 mouse telencephalon. **(F)** Motif and evolutionary conservation landscape for enhancers in (d) showing motif clustering and overlap with conserved regions across core 500bp (top) and at single-base resolution (bottom) for selected intervals. Legend shows colors for BG TF primary motifs and all Homer motifs. See also Figure S5.

We next examined spacing and orientation of motif pairs within pREs across clusters. Overall, cluster-relevant motif pairs were likely to be closely spaced (i.e., within 20-200 bp distance) within pREs (Figure 5B). This is most obvious for pRE clusters with more motif instances. For example, TAATTA-TAATTA pairs were located in spatial proximity within pRE clusters featuring broad TF binding, epitomized by 1_D. 1_D also exhibited reduced but still clear proximity between TAATTA-NKX2.1 and TAATTA-PBX1 pairs, though not for TAATTA-ASCL1. In comparison, 12_D, bound specifically by ASCL1, showed clustered ASCL1-ASCL1, but not TAATTA-TAATTA, pairs. There were also differences in TAATTA-TAATTA pairs across clusters. For example, 4_D and 16_D both feature similar average number of TAATTA instances, but TAATTA-TAATTA pairs are more likely immediately adjacent in 16_D, with 26% of pREs featuring a pair of TAATTA motifs within 4 bp compared to 8% in 4_D and 10% in 1_D. We did not identify canonical orientation or spacing rules for motif pairs in any of the TF-pRE clusters, as would be predicted if TF complex binding was determined by a “syntax” of motifs with deterministic spacing. At the upper end of the spectrum for motif density, 1_D featured clustered, and at times overlapping, sets of motifs including the primary motifs of TFs studied here as well as other motifs in the HOMER database. Overall, our findings are consistent with a “billboard” model, where relevant DNA binding sequences are clustered within a core region of the regulatory DNA element, but do not conform to strict patterns of orientation or syntax^36^.

Lastly, we examined base-level PhyloP vertebrate conservation of TF motifs within pREs (Figure 5C). Overall, motifs expected to be relevant for TF binding showed increased base-level conservation in line with motif position weight matrix and reduced relative conservation for flanking sequences, indicating purifying selection on bases critical for TF binding (see Figure S5a for complete motif by cluster analysis). For example, average base-level conservation for DLX2 primary motif shows strong conservation for the TAATTA bases compared to flanking sequence for both cluster 1_D and 16_D. Base-level motif conservation also showed differences across pRE clusters bound by the TF. While background conservation is significantly higher in 1_D versus 16_D, the TAATTA core also shows a larger increase relative to background in 1_D. For DLX2 TAATTA motifs in 1_D, increased base-level conservation of flanking DNA gradually decreases out to +/-200bp, indicating a conserved core region within these pREs. We next expanded base-level motif conservation analysis to all motifs in the Homer database that were enriched in each pRE cluster. Considering the evidence for the central role of TAATTA motifs in 1_D, we wondered if similarly strong base-level conservation was present across other motifs. Motifs from several TF families indeed exhibited increased conservation, for example HOXD11/Hox, FOXA1/Forkhead, OCT6/OCT, and SOX6/SOX (Figure 5D). Among motifs with high base-level conservation in 1_D, TAATTA motifs are at the top of the range, cementing a special role for TAATTA motifs and TFs that bind these sequences in 1_D enhancers.

Our results indicate that at the sequence level, 1_D pREs are characterized by increased number of symmetric TAATTA motifs, which are located in proximity to each other and to other motifs in a highly-conserved and motif-rich core region. To illustrate these patterns, four representative newly-characterized 1_D pREs are depicted in Figure 5E. These 1_D enhancers capture the overall patterns of this cluster, giving examples of the motif clustering but variability of spacing and organization as well as the base-level motif conservation both across the enhancer core as well as of critical nucleotides within each motif. These results provide evidence that enhancers at the extreme upper end of evolutionary conservation indeed feature dense hubs of cis-trans regulatory interactions during embryonic development that are likely to drive strong selective constraint.

Our results highlight the complex regulomes of neurodevelopmental TFs targeted by the TFs profiled here, potentially associating distinct genes via common chromatin interaction ensembles. Actively regulated chromosomal regions, particularly those harboring TF genes, feature complex physical interaction landscapes anchored by differential combinations of TFs that act to activate or repress enhancer activity in the developing BG. As an example, for the interaction ensemble including the *Pbx1* gene, we identified 31 pREs that serve as PSCs (Figure 6A) and transgenic testing verified activity for three enhancers predicted to be activated in BG and corresponding pallial or non-CNS activity for enhancers correctly predicted to be repressed in BG (Figure 6B).

**Figure 6.**
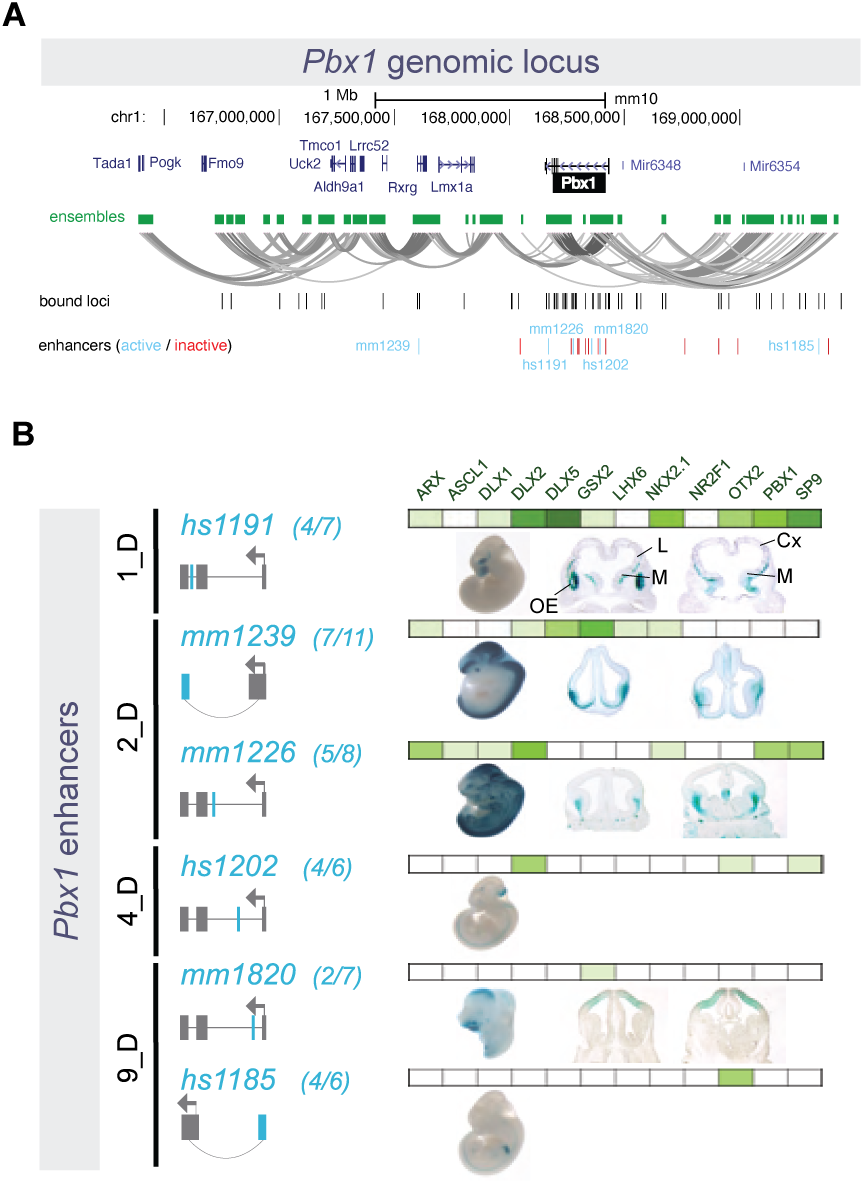
*Pbx1* Genomic Locus and Associated Putative Enhancers. **(A)** Representation of the *Pbx1* locus, showing nearby genes, and the ensemble of PLAC-seq contacts creating a tridimensional structure made of multiple loops. Bound genomic loci are noted underneath in black, and enhancers are marked in turquoise and red (active and inactive, respectively). **(B)** Six enhancers around the *Pbx1* locus exhibited activity in transgenic mouse assays. Of these, 1_D and 2_D enhancers were active in subpallium, while one 9_D enhancer was active in in pallium and the other 9_D and a 4_D enhancer showed non-telencephalic activity.

## DISCUSSION

Application of epigenomic profiling resolved to cell type specificity have revolutionized understanding of cis-regulatory landscapes underlying neurodevelopment, including in the embryonic BG^37^. Missing from these advances has been parallel comprehensive understanding of TF components of gene regulatory wiring, and, specifically, how combinatorial TF binding to cREs directs transcriptional activation and repression. Studies focused on individual or a small set of TFs have provided valuable insights about necessity and sufficiency of specific TFs during neurodevelopment and in the BG^1^ and cortex^38^. Yet an integrated perspective of TF binding during brain development has been lacking due to technical and computational barriers, particularly for ChIP-seq on *ex vivo* embryonic brain tissues. *In silico* and *in vitro* efforts to use TF co-expression and motif analyses have provided insights into contributions of TF networks even without assaying genomic interactions^39–41^. However, modeling roles of TFs without interrogating genomic binding is problematic due to shared target motifs and overlapping expression within homeobox and other TF families, and due to presence of indirect TF interactions with regulatory DNA. Further complicating functional modeling, our results corroborate previous understanding that not all TF binding events are equivalent regarding regulatory function^42, 43^. Illustrating these complexities, all tested TFs (except for NR2F1) participated in more than one distinct combinatorial TF binding pattern that had different functional roles, six of the homeobox TFs bound the same family of TAATTA motifs, SP9 primarily bound cREs via indirect mechanisms. This study revealed patterns of TF-cRE interaction and regulatory function that would be masked if only considering chromatin accessibility or individual TF binding patterns. This work represents an initial build of an embryonic BG “regulome” made up of TF-cRE modules that integrates information for TF binding, chromosomal interactions, chromatin state and transcriptional activity, and cRE regulatory function.

A motivating factor for this study was to define complex patterns of the combinatorial TF binding in the developing brain, going beyond previous work on individual TFs to build models similar to efforts in other organisms and for in vitro systems^50–53^. We had previously found that DLX TFs had a largely overlapping set of binding targets, and NKX2-1 and LHX6 had overlapping and distinct binding targets^18, 20^. Here we combined published and novel TF ChIP-seq across twelve TFs, capturing diversity of TF families and neurodevelopmental function. Our work illustrates context-dependent combinatorial TF control of activating and repressive cRE modules and gene regulatory programs across proliferating and post-mitotic neuronal populations in embryonic mouse BG (Figure 7A). We show that TF-cRE interactions are required for proper activation and repression of enhancer activity associated with sub-pallial regional identity, GABAergic neurogenesis, and cortical interneuron specification (Figure 7B). These different TF-cRE modules act in parallel to generating complex developmental expression patterns, for example at the *Pbx1* locus (Figure 7C). These results reveal the molecular and genomic basis underlying findings from earlier TF knockout mouse studies for *Gsx2, Nkx2-1,* and *Otx2* showing shifts in telencephalic regional identities^21, 27, 44^, and for *Arx, Ascl1, Dlx1/2/5, Nr2f1, Pbx1,* and *Sp9* showing decreased GABAergic neurogenesis and differentiation^19, 22, 45–49^. Our integrated characterization of embryonic BG TF-pRE modules and the 74 newly-defined enhancers represent a rich resource for future studies of transcriptional control of enhancer function, enhancer-driven cell labeling, and neuronal fate mapping.

**Figure 7.**
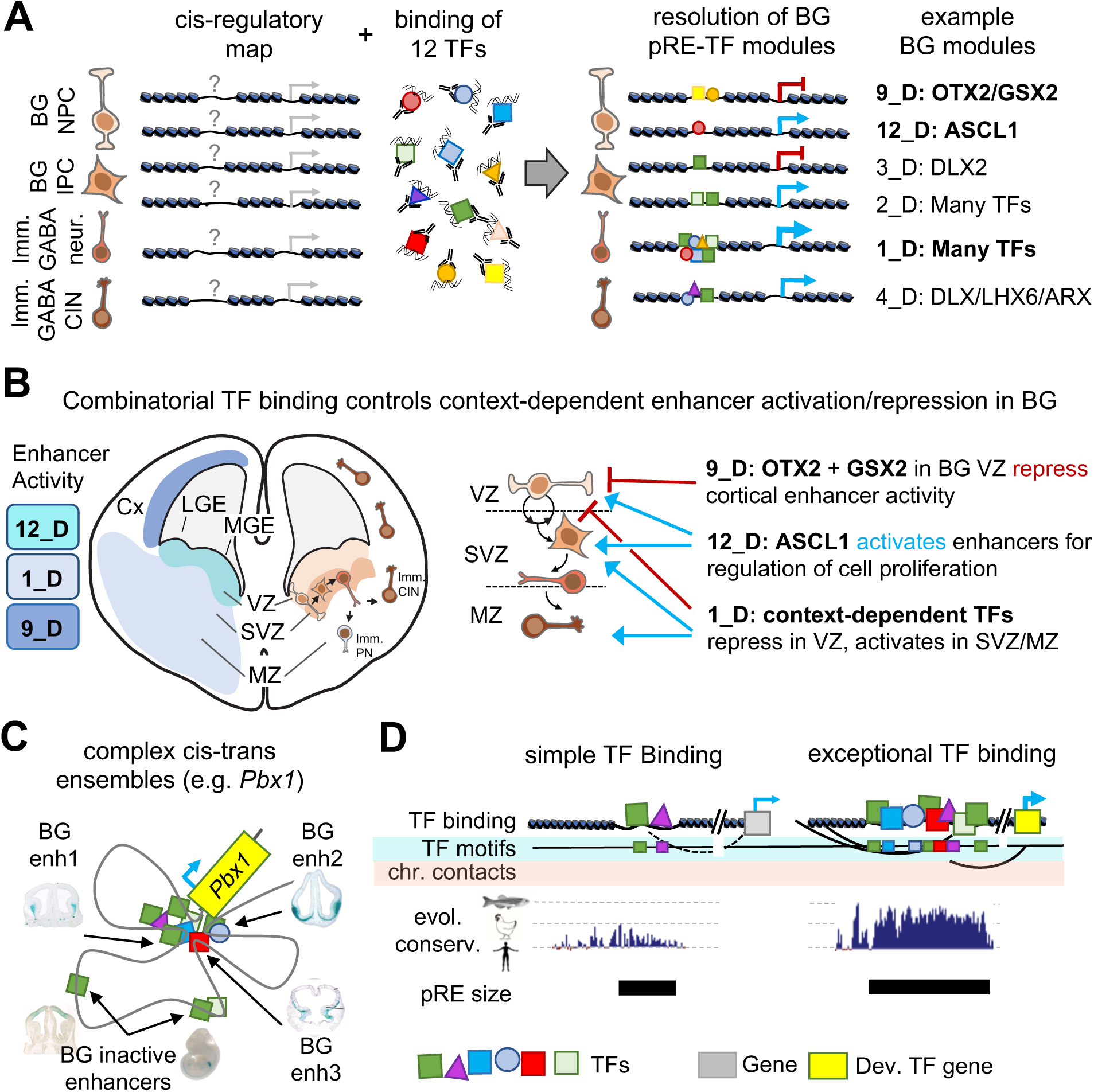
Cis-trans interactions underlying gene regulation driving GABAergic neurogenesis. **(A)** Chromatin accessibility maps identify pREs, but TF binding is necessary to understand mechanisms and functional relevance of pRE activity. TF binding can direct either activation or repression of enhancer activity. Here we identify pRE-TF modules that drive specific regulatory activity in developing mouse BG, with representative examples depicted. Bold BG modules in (A) are highlighted in (B). **(B)** Combinatorial TF binding defines context-dependent patterns of enhancer activation and repression in embryonic BG. Three example cis-trans modules identified here are shown, with the enhancer activity and schematic of activity across VZ, SVZ, and MZ. **(C)** Developmental TF genes (i.e. *Pbx1)* relevant to embryonic BG have complex cis-regulatory landscapes and generally include multiple cis-trans regulatory modules. **(D)** Comparison of enhancers with simple versus complex TF binding identified in embryonic BG. Enhancers with exceptional TF binding also feature high density of TF binding motifs, complex chromosomal contacts, strong evolutionary conservation across the vertebrate tree (human, chicken, zebrafish conservation represented), and increased base pair size. Abbreviations: Imm. CIN: immature cortical interneurons, Imm. PN: immature projection neurons, BG NPC: basal ganglia neural progenitor cell, BG IPC: basal ganglia intermediate progenitor cell.

Among the most important insights from our analysis was the overlap and strength of TF binding across 11 of the 12 TFs at a relatively small set of distal pREs. These findings support previous studies that TF binding overlap is associated with increased enhancer activity and conservation^4, 54, 55^. For example, pREs with OCT4, SOX2, and NANOG combined binding had stronger evolutionary conservation and enhancer function in ESCs than sites with single TF binding^56^. Indeed, 1_D included an enrichment for elements that were identified as among the most conserved non-coding sequences in the mouse genome^57^. Our findings build context around observations made regarding what have been deemed “ultraconserved” enhancers, including that these enhancers are key partners for developmental TFs, are strongly enriched for activity in the developing brain, that they are enriched for TA-rich motifs and robustly bound in vivo by homeobox TFs in this study^58–60^, but whose defects are nonetheless non-lethal^61^. To this picture, our results show that BG-active deeply-conserved enhancers are broadly bound by TFs, feature rich and often overlapping motif composition and extreme base-level conservation (Figure 7D). Our results show that homeobox TFs, and TAATTA-binding TFs in particular, bind this set of ancestral enhancers, which regulate genes that act at the top of signaling pathways that regulate the regulators of brain development^62, 63^. Our study further supports the model where these extensive TF-enhancer interactions are involved in establishing and maintaining complex transcription-associated local biophysical interactions^64^. Our results frame a model where cREs that are broadly TF bound during neurodevelopment represent the extreme end of the spectrum for sequence composition, conservation, and regulatory activity, and as such are exceptional and ancient class of enhancers.

## AUTHOR CONTRIBUTIONS

Conceptualization, R.C.-P., A.S.N., S.L., and J.L.R.R.; Methodology and Investigation, S.L. (ChIP-seq, RNA-seq), Y.S. (PLAC-seq experimental), A.A., M.H. (PLAC-seq computational), J.D.P. and L.S.-F. (RNA-seq), D.E.D., L.A.P., and A.V. (enhancers in mice), and R.C.-P. and Y.W. (bioinformatics, modeling); Software, R.C.-P., A.S.N., and A.A.; Formal Analysis, R.C.-P., A.S.N., S.L., and J.L.R.R.; Writing – Original Draft, R.C.-P., S.L., A.S.N., and J.L.R.R.; Writing – Review & Editing, R.C.-P., S.L., J.D.P., D.E.D., A.S.N., and J.L.R.R.; Funding Acquisition, J.L.R.R. and A.S.N.; Supervision, J.L.R.R. and A.S.N.

## ACKNOWLEDGEMENTS

University of California, San Francisco (UCSF) and Davis (UCD) cores supported this work. Sequencing was carried out at the UCSF Center for Advanced Technology and IHG Genomics Core, UCD DNA Core. R.C.-P. was supported by a Science Without Borders Fellowship (Conselho Nacional de Desenvolvimento Cientifico e Tecnologico - CNPq, Brazil). L.S.-F. was supported by the UCD Schwall Fellowship in Medical Research and Werner and Jacobsen Fellowship, and National Institute of General Medical Sciences (NIGMS)-NIH T32-GM008799. This work was supported by the following research grants: National Institute of Mental Health (NIMH) R01 MH081880 and NIMH R37/R01 MH049428 (to J.LR.R.); NIH/NIGMS R35 GM119831 (to A.S.N.); NIH U01DA052713 (to Y.S.). A.V., L.A.P., and D.E.D. were supported by NIH grants R01MH117106, R01HG003988, and R01NS062859. Research was conducted at the E.O. Lawrence Berkeley National Laboratory and performed under Department of Energy Contract DE-AC02-05CH11231, University of California.

## SUPPLEMENTARY MATERIAL

Supplementary Tables S1-S6

Supplementary Figure S1 (Related to Figure 1)

Supplementary Figure S2 (Related to Figure 2)

Supplementary Figure S3 (Related to Figure 3)

Supplementary Figure S4 (Related to Figure 4)

Supplementary Figure S5 (Related to Figure 5)

## MATERIAL AND METHODS

### Experimental Model and Subject Details

#### Mice

All procedures and animal care were approved and performed in accordance with National Institutes of Health and the University of California San Francisco Laboratory Animal Research Center (LARC) guidelines. For the RNA-seq experiment, an equal number of males and females were used. ChIP-seq and native histone ChIP-seq was performed on *Mus musculus* CD1 strain at developmental stage E13.5. The embryos were not assessed genotypically for gender since we used a pool of embryos and therefore expect a roughly equal number of male and females.

### Method Details

#### TF Chromatin Immunoprecipitation (ChIP)

ChIP was performed using antibodies against DLX1, DLX2, DLX5^18^, NKX2.1 (Santa Cruz Biotechnology, Cat# sc-13040), LHX6^20^, OTX2 (Published in Hoch, Lindtner, Price and Rubenstein - R&D, Cat# AF1979), SP9^65^, ARX (Santa Cruz, Cat# sc-48843), ASCL1 (BD, Cat# 556604), GSX2^66^, NR2F1 (R&D biosystems, Cat# PP-H8132-10), and PBX1/2/3 (Santa Cruz, Cat# sc-888). Basal ganglia were dissected in cold PBS from CD1 embryos (2 L/Ab for DLX2, SP9, NR2F1; 3 L/Ab for DLX1, DLX5, NKX2.1, LHX6, ARX, ASCL1, and GSX2; 2 L/Ab for OTX2). The basal ganglia consisted of the LGE, MGE and CGE progenitor and mantle zones except the NKX2.1 and LHX6 ChIPs for which the medial ganglia were used. The dissected basal ganglia were either fixed in 1% formaldehyde at RT for 10 min (LHX6, NKX2.1, OTX2, PBX, SP9) or fixed in 1.5% formaldehyde at RT for 20 min (ARX, ASCL1, DLX1, DLX2, DLX5, GSX2, NR2F1, OTX2), neutralized with glycine, and washed gently in PBS. The fixed cells were lysed with a hypotonic buffer (50 mM Tris pH 7.5 / 0.5% NP40 / 0.25% sodium deoxycholate / 0.1% SDS / 150 mM NaCl) to obtain the nuclei; these were then lysed in 1% SDS buffer and the chromatin was sheared into 300-1000 bp fragments by sonicating for 40 cycles (30 sec on and 45 sec off) using a bioruptor (Diagenode). Immunoprecipitation (IP) reactions were performed with the sheared chromatin diluted 1/10 times with “dilution buffer” (0.01% SDS, 1.1% Triton X-100, 1.2 mM EDTA, 16.7 mM Tris-HCl, pH 8.1, 167mM NaCl, usually in 6 ml. Antibody was then added to either 5 µg (ARX, DLX1, DLX2, DLX5, NR2F1, OTX2, PBX, SP9) or 8 µg (ACSL1, GSX2, LHX6, NKX2.1) specific antibodies. Negative control ChIP reactions used either IgG (5µg) or blocking peptide (DLX antigen used for immunizing rabbits; 50x molar excess, ARX, LHX6, NKX2.1, PBX 400x molar excess). Antibody/chromatin complexes were purified using Dynabeads (Invitrogen) and washed extensively in “wash buffer” (low salt, 0.1% SDS, 1% Triton X-100, 2 mM EDTA, 20 mM Tris-HCl, pH 8.1, 150 mM NaCl; high salt, 0.1% SDS, 1% Triton X-100, 2 mM EDTA, 20 mM Tris-HCl, pH 8.1, 500 mM NaCl; LiCl, 0.25 M LiCl, 1% IGEPAL CA630, 1% deoxycholic acid (sodium salt), 1mM EDTA, 10mM Tris, pH 8.1 and TE).

Complexes were eluted with 1% SDS, 10 mM sodium bicarbonate buffer at 65 °C for 10 min. Eluted chromatin was reverse-crosslinked overnight at 65 °C in the presence of 500 mM NaCl, then subsequently treated with RNase (10 µg/200 µL reaction, 15 min at 37 °C) and Proteinase K (10 µg/200 µL reaction, 60 min at 55 °C) and cleaned using a ChIP DNA Clean & Concentrator kit (Zymo Research). The chromatin was quality-controlled (QC) using qPCR to check for enrichment of genomic DNA fragments that were expected, and not expected, to have the different TF binding.

Libraries were prepared using an Ovation Ultralow DR Multiplex System (Nugen), size-selected in the range of 300 bp on a chip from BluePippin (Sage Science) and lastly QC tested on a Bioanalyzer (Agilent). The libraries were sequenced as single-end 50-bp reads on a HiSeq 4000 (Illumina) at the Center for Advanced Technology (UCSF).

#### Native histone ChIP

Each native histone ChIP was performed starting with ∼250,000 nuclei from WT E13.5 basal ganglia. The native ChIP was performed as described earlier^18^. Briefly, nuclei were extracted and digested with micrococcal nuclease (MNase, Sigma). A population of mono-and di-nucleosomes were used in chromatin immunoprecipitation assays. Antibodies used were specific to H3 monomethyl lysine-4 (H3K4me1, Abcam, ab8895), H3 trimethyl lysine-4 (H3K4me3, Abcam, ab8580), H3 trimethyl lysine-27 (H3K27me3, Active Motif, 39157), and H3 acetylated lysine 27 (H3K27ac, Abcam, ab472). Immunoprecipitated DNA was washed, isolated, and cleaned as for the TF ChIP-seq described above.

#### PLAC-seq

PLAC-seq libraries for E13.5 basal ganglia were prepared similar to a previously published protocol^67^. 3 to 7 million cells were used for each library. If the cells appeared aggregated, they were dissociated with gentle MACS dissociator or Dounce homogenizer. Each PLAC-seq library was prepared using *DpnII* as the restriction enzyme and Dynabeads M-280 sheep anti-rabbit IgG (Invitrogen #11203D) mixed with 5 µg of H3K4me3 (04-745, Millipore) for the chromatin immunoprecipitation step. Finally, libraries were prepared with the Illumina TruSeq adaptors and the final libraries were sent for paired-end sequencing on the HiSeq X Ten (150-bp reads) equipment.

#### Transgenic enhancer assays

All transgenic enhancer assays were performed at Lawrence Berkeley National Laboratory (LBNL) under the approval of the Animal Welfare and Research Committee (AWRC), as previously described^18^. In short, candidate enhancers were PCR-amplified and cloned into an *hsp68*-promoter-*lacZ* reporter vector^68^. Transgenic assays were performed according to published methods^68, 69^. The enhancer-reporter vector was linearized and injected into the pronucleus of FVB strain single cell stage mouse embryos (E0.5). Embryos were implanted into surrogate CD-1 strain *Mus musculus* mothers and were then collected and stained for reporter gene expression at E11.5, E12.5, or E13.5. The resulting embryos were not assessed phenotypically for gender, which is not outwardly obvious at these ages. Therefore, we expect that a roughly equal number of male and female embryos were assessed. Embryos were excluded from analysis only if they did not harbor the transgene or if they were not at the correct developmental stage. No comparisons were made between cohorts of transgenic embryos, so randomization and experimenter blinding were unnecessary and not performed. Sample sizes were determined empirically based on our experience performing >2,000 transgenic enhancer assays. Only *LacZ* activity patterns that were observed in more than 30% of embryos resulting from independent transgene integration events of the same construct were considered reproducible.

#### Histology and Regional Activity Scoring

Brains were fixed, cryopreserved and embedded as described previously^70^. *LacZ* activity of candidate enhancers were annotated by at least two experts in the field by annotating activity in the ventricular zone and the subventricular zone/mantle zone of the ganglionic eminences.

### Computational and Statistical Analysis

#### PLAC-seq

We detected the statistically significant long-range chromatin interactions from H3K4me3-associated proximity (PLAC-seq) data, using the MAPS pipeline^71^. Only intra-chromosomal interactions for autosomal chromosomes were selected, at a 10-kb resolution in the range 20 kb to 1 Mb. Raw reads from sequencing (FASTQ files) were mapped to the mm10 genome annotation using BWA mem. After duplicate, chimeric and low-quality read removal^71^, we split the mapped reads into short-and long-range reads, for distances between pair ends less than 1 kb and in the 1 kb-1 Mb range, respectively. We used the short-range and long-range reads to measure protein immunoprecipitation (IP) efficiency and detect long-range chromatin interactions, respectively.

For ascertaining meaningful interactions, we called peaks from ChIP-seq experiments on GE cells using H3K4me3 antibody using MACS2^72^. Only 10-kb bin pairs for which at least one end overlapped with called ChIP-seq peaks were used in the analysis. We fitted a positive Poisson regression model on all selected 10-kb bin pairs with more than one raw count, taking into consideration bias factors including linear genomic distance between two interacting bins, restriction enzyme cut site frequency, GC content, mappability score, and H3K4me3 antibody efficiency measured by the number of short-range reads in each bin. After model-fitting, we obtained expected contact frequency, p-value and false discovery rate (FDR) for each 10-kb bin pairs. We filtered only statistically significant bin pairs, defined as those with (1) raw contact frequency >= 12, (2) normalized contact frequency (observed/expected contact frequencies) >= 2, and (3) FDR < 0.01.

We defined singletons as isolated significant chromatin interactions that passed the very stringent FDR of 1 x 10^-4^ (to reduce potential false positives) after merging adjacent chromatin interactions together. Contiguous contact point intervals from the table generated as above were merged, and the resulting interaction graphs were produced and analyzed under the denomination *interaction ensembles*, using a custom R script.

#### RNA-seq Data Analysis

Gene expression in GEs at E13.5 were expressed as log_2_(RPKM) and calculated from the mean of read counts assigned to genes using the UCSC annotation. RNA-seq data was generated as part of a study previously reported^18^, and was used without further modifications.

#### ChIP-seq Data Analysis

The basic analysis pipeline is depicted in the Figure S1a. Quality-controlled FASTQ files^73^ containing the reads were further cleaned-up from the remaining adapter sequences using Trim Galore version 0.4.5^74^. The resulting reads were aligned to the mouse (mm10) genome using BWA version 0.7.9a^75^, and duplicates removed with Samtools version 1.8^76^. Peaks of enriched binding regions against both negative binding and input DNA control were called using MACS version 2.1^72^ with p-value cutoffs of 1 x 10^-4^.

To infer co-occurrence of TF binding at each genomic locus, we combined the peaks called from 6 published and 6 novel TF ChIP-seq datasets, as described above. Narrow peaks of TFs were overlapped and merged into one dataset using custom R scripts, where they were annotated for neighboring gene regulatory regions and filtered against repeat, blacklisted, and gapped regions. For each TF with more than one replicate, we selected one that was representative of the whole set; we ran Pearson correlation analysis on genome wide coverage using DeepTools version 2.5.3^25^ and determined that replicates for the same TF were consistent.

Hypothesizing that genomic context of binding is determined not only by the binding loci but also by their neighborhood, we used DeepTools to cluster read coverage for each TF around 1 kb (each side) of each peak in the combined data set at 10-bp resolution employing the K-means strategy. The data set was split into proximal and distal regions, defined as those either overlapping or not with putative gene promoters (respectively 2000 and 200 bp upstream and downstream of TSSs), respectively. The number of clusters was initially set to 3 for each distal/proximal subset and increased by one until no further patterns were visually captured by the authors, except slight variations in the clusters bound by multiple TFs. We visually assessed the clusters to determine patterns. In the distal subset, one of the clusters encompassed more than 8000 peaks; we further clustered that cluster using the same strategy, and found additional patterns as outlined above. In the proximal subset, two clusters had similar binding profiles, and one of them was too small to allow statistical analysis; they were combined, and shown in Figure 2b as 2.1_P and 2.2_P.

#### Motif Analysis

Core motifs for each of the 12 TFs were determined using HOMER version 4.9^77^ in the called peaks for each individual TF. We performed *de novo* motif discovery with standard parameters, 300 bp up-and downstream of TF peaks. We compared the highest significant discovered motifs with HOMER known motif and JASPAR^78^ databases, and inferred the core motif based on similitude. For all TFs except SP9 the core motif was the most significant one (lowest p-value from the HOMER analysis). The motif enrichment in the combined data set was determined for each cluster, distal and proximal loci combined, and overall, for all motifs present in the HOMER known motif database with p-value < 10^-125^ and enrichment > 1.5 (unless otherwise noted). We removed from the analysis all motifs from proteins encoded by genes not expressed in E13.5 GEs, as determined by the RNA-seq experiment.

For each individual TF and bound loci cluster, the average motif coverage enrichment plot around 300 bp of peaks was established using HOMER annotation output and custom R scripts. For the coverage analysis on clustered bound loci, we averaged the signals from the homeobox-containing motifs among our 12 TFs. The heatmap in Figure 2f reflects individual core motif prevalence, whose relative enrichment is displayed in Figures S2a and S2b, under different scopes. The average core motif coverage plots across the different binding clusters are shown in Figure S2d.

#### Integration of orthogonal data types

To make the inferences described in the text, we integrated transcriptomic (RNA-seq), binding (ChIP-seq), epigenomic (histone ChIP-seq), and interactomic (PLAC-seq) data, using custom R scripts (Figure S1a). We overlapped binding loci with chromatin interactions (both ensembles and single interactions), and histone (H3K27ac, H3K27me, H3K4me3, and H3K4me1) coverage across the mm10 genome. Those intersection loci were further stratified into the neighborhood binding clusters, as laid out above. We also overlapped those stratified loci against transcriptional enhancers from the VISTA database^32^, as well as novel ones generated as part of this study.

Additionally, gene-ensemble association was annotated, based on the presence of annotated gene promoters in interaction ensembles. Those interactions were used to infer a gene expression model from the transcriptional regulation by three-dimensional chromatin combinatorially-bound by the TFs. In doing so, we also employed transcriptomic data in published mouse E13.5 GEs^18^.

#### Chromatin State Inference

We created a 9-state HMM model of chromatin states using E13.5 BG WT histone data previously reported^18^ and ChromHMM^79^ and assigned chromatin states based on the emission probabilities. The number of states was determined by the minimum time for convergence of the algorithm as well as assessment for biological relevance. Symbols were assigned to each state parallel to previously published assessment^80^. Our model chromatin states were assessed as below.

| HMM state | Assigned symbol | Chromatin state |
| --- | --- | --- |
| 1 | EnhLo | active enhancer with transcriptional history |
| 2 | EnhTx | highly active enhancer with transcriptional history |
| 3 | Enh | active enhancer without transcriptional history |
| 4 | EnhPoi | poised/weak enhancer |
| 5 | EnhBiv | bivalent enhancer with transcriptional history |
| 6 | ReprPC | polycomb-repressed locus |
| 7 | Quies | no signal (quiescent) |
| 8 | TxWk | weak transcriptional signal |
| 9 | TssBiv | bivalent promoter element |

#### Other inferences

Non-Gaussian distribution comparisons were performed by computing sample means (N = 10000) and comparing the means distributions by pairwise t-test; samples were collected without replacement with a size of 75-80% of the sampling space. For the log likelihood comparisons between observed and expected, the expected value was derived from the frequency of the category in each class as per the overall frequency of that category (Figure S3).

We compared our results with published data to validate our inferences. We intersected our neighborhood binding clusters with the clusters derived from the analysis of single-nucleus chromatin accessibility (scATAC-seq) of developing and post-natal mouse forebrain^28^, using custom R scripts. Predominant cell fates in those clusters emerged from that overlap, allowing the inference of TF roles in the regulation of forebrain development in the GEs.

For the assessment of overall and base-level evolutionary conservation of stratified genomic loci as previously described, we used published Vertebrate Phastcons (60 species)^81^ and PhyloP (59-way)^82^ scores for the mm10 mouse genome, respectively, downloaded from the UCSC genome browser (http://hgdownload.cse.ucsc.edu/goldenPath/mm10/).

We further validated our model by comparing the inferences with previously published findings of regulatory roles of DLX2^18^ and NKX2.1^20^ in activating (a.RE) and repressing (r.RE) gene expression.

#### Comparison with Cortex TF binding

To make the comparison against known TF binding in developing cortex at a similar developmental stage, we reanalyzed published cortex ChIP-seq datasets^38^ under the same parameters herein, and overlapped those binding loci with those obtained in our study, using custom R scripts.

#### Data and Code Availability

This article made use of published and unpublished genomic and epigenomic data that are deposited in the NCBI database, as outlined below. The data can be downloaded from (https://www.ncbi.nlm.nih.gov/geo/query/acc.cgi?acc=GEO *accession*), as below.

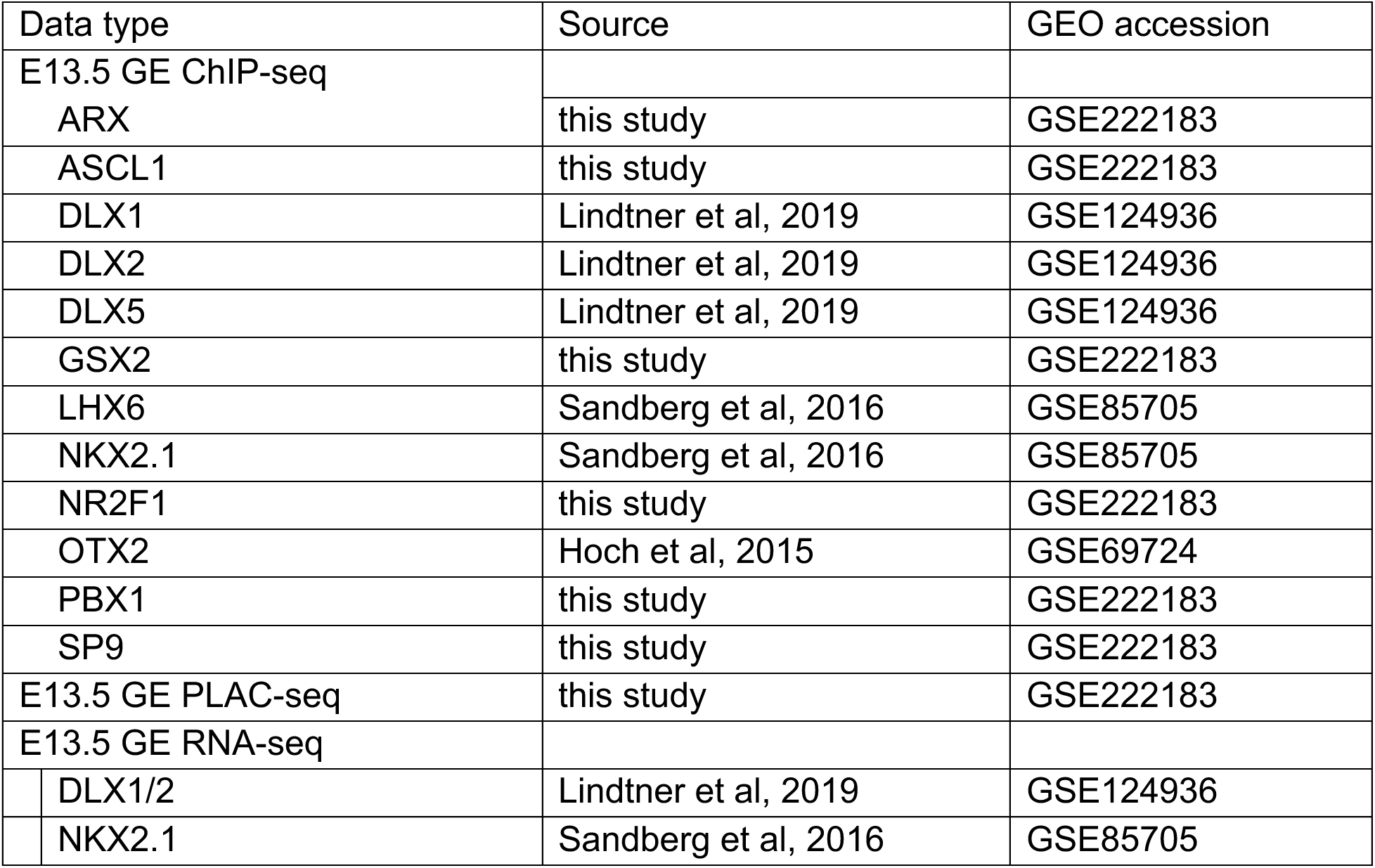

Data can be visualized in UCSC track hub format whose information is provided on Nord Lab GitHub page (https://github.com/NordNeurogenomicsLab/Publications/tree/master/Catta-Preta_XXX_2023). All R scripts were created in-house and can be provided upon request.

