## Supplementary Fig 1 for "Combinatorial transcription factor binding encodes cis-regulatory wiring of forebrain GABAergic neurogenesis"

### Supplementary Figure S1

**TF binding, chromosomal interactions, and chromatin state in E13.5 mouse basal ganglia. (a)** Schematic of workflow for computational analysis pipeline used in this study. Dark green boxes are the sources of raw data and blue boxes contain the unit processes leading to the integration. Arrows/lines indicate the sequence of analysis. Under each box the software, utility or algorithm used to perform the operation is noted. See **Methods** for details. **(b)** *In situ* hybridization assays showing regional RNA expression of tested TFs in embryonic mouse brain at E13.5 (https://developingmouse.brain-map.org/). Cx: Cortex, LGE: Lateral Ganglionic Eminece, MGE: Medial Ganglionic Eminence, Se: Septum, D: Diencephalon, Ch Pl: Choroid Plexus, VZ: Ventricular Zone, SVZ: Sub-ventricular Zone, MZ: Mantle Zone. **(c)** Single cell RNA-seq (snRNA-seq) data showing expression of 12 TFs from analysis of E12.5 mouse BG. Each point shows expression in a single cell. Y axis is relative transcript expression and x axis is developmental pseudotime generated by diffusion mapping analysis. Data previously published in [1]. Left side of the x-axis represents earliest progenitor populations in ventricular zone and right side represents most mature post-mitotic neurons in mantle zone. **(d)** Boxplot showing mean loci width distribution relative to the number of co-binding TFs across distal and proximal regions. General relationship of increasing locus width as more TFs bind to the locus is due to merging of partially overlapping TF ChIP-seq peaks as well as wider ChIP-seq signal for individual TFs at loci featuring extensive binding. **(e)** Bar plots showing percent overlap of bound TF peaks and PSCs for proximal and distal loci for each TF profiled. **(f)** Same as (e), but for proportion of TF-bound loci within chromatin loops defined by the PSCs. **(g)** Histogram showing the distribution of “complexity” of PLAC-seq defined chromatin interaction ensembles, where complexity is measured by the number of interacting PSCs. **(h)** Cumulative distribution plot, depicting the percent of ensembles bound by at least 1 TF as a function of ensemble size, showing that after a relatively small ensemble size, all ensembles feature at least one PSC bound by combinations of the 12 TFs profiled here. **(i)** Heatmap showing the proportion of chromatin state for TF-bound loci annotated to PSCs, and inside loops (regions between PSCs), or outside of loops. PSCs are annotated to transcribed and active enhancers and bivalent TSSs, as expected based on use of H3K4me3 antibody for PLAC-seq pulldown. **(j)** Heatmap showing the proportion of chromatin states by genome occupancy, colocalization with CpG islands, overlap with gene exons, gene bodies, TES (transcription end sites), at or within 2kb of TSS (transcription start sites). Also included are the emission probabilities for the 4 histones PTMs at each state and annotation by activity. The bars to the right of the plot portraits the interpretation of each state (EnhLo = active enhancer with transcriptional history; EnhTx = highly active enhancer with transcriptional history; Enh = active enhancer without transcriptional history; EnhPoi = poised/weak enhancer; EnhBiv

= bivalent enhancer with transcriptional history; ReprPC = Polycomb-repressed locus; Quies = no signal or quiescent).


1. Su-Feher, L. *et al.* Single cell enhancer activity distinguishes GABAergic and cholinergic lineages in embryonic mouse basal ganglia. *Proc. Natl. Acad. Sci.* **119**, e2108760119 (2022).

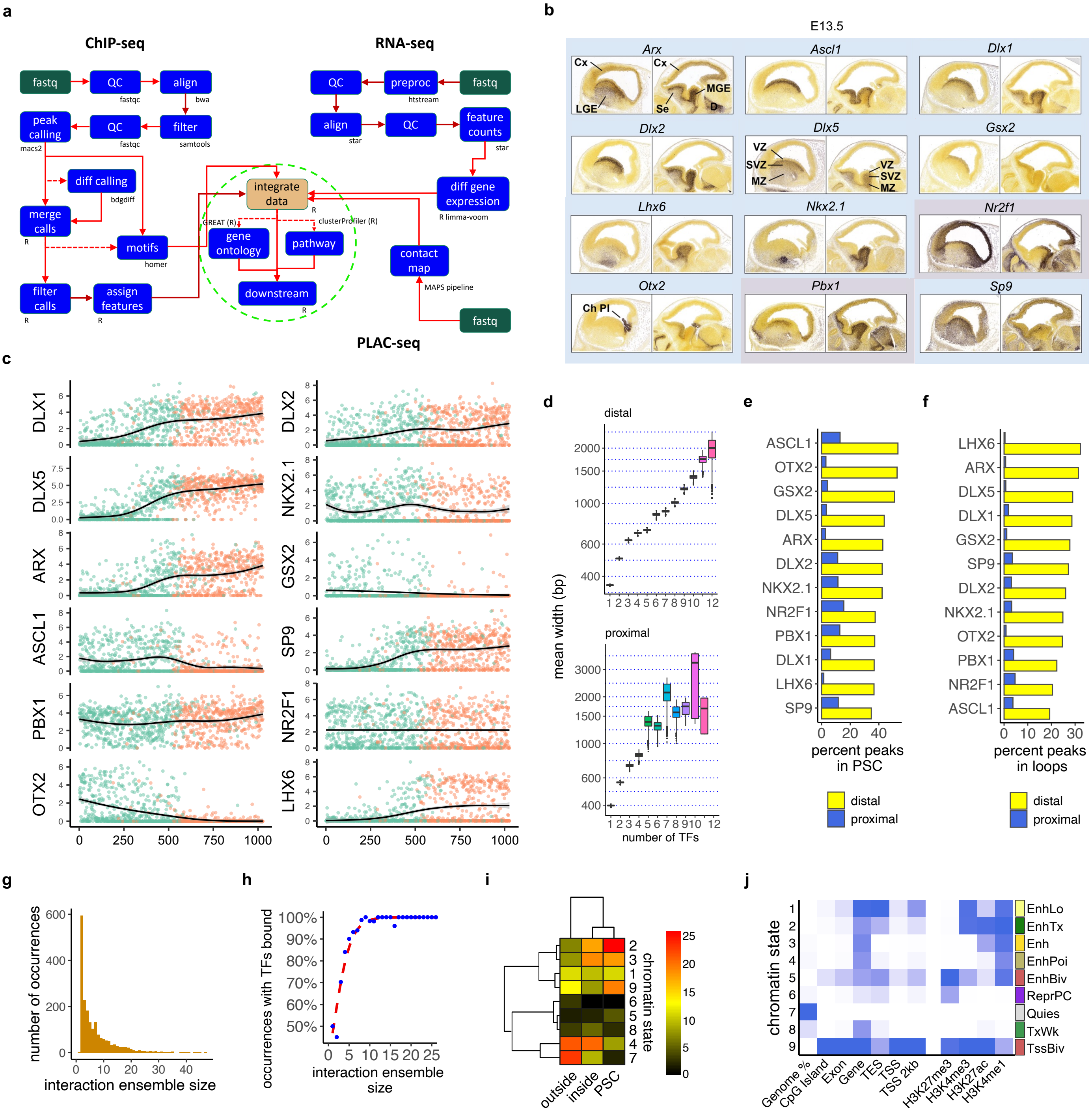
a
ChIP-seq
RNA-seq
PLAC-seq
fastq
QC
fastqc
align
bwa
peak calling
macs2
QC
fastqc
filter
samtools
diff calling
bdgdiff
merge calls
R
motifs
homer
filter calls
R
assign features
R
integrate data
R
GREAT (R)
gene ontology
clusterProfiler (R)
pathway
downstream
R
QC
preproc
htstream
fastq
align
star
QC
feature counts
star
diff gene expression
R limma-voom
contact map
MAPS pipeline
fastq
b
E13.5
Arx
Cx
Cx
MGE
LGE
Se
D
Ascl1
Dlx1
Dlx2
Dlx5
VZ
SVZ
MZ
VZ
SVZ
MZ
Gsx2
Lhx6
Nkx2.1
Nr2f1
Otx2
Ch Pl
Pbx1
Sp9
c
DLX1
DLX2
DLX5
NKX2.1
ARX
GSX2
ASCL1
SP9
PBX1
NR2F1
OTX2
LHX6
0
250
500
750
1000
d
distal
proximal
mean width (bp)
number of TFs
e
ASCL1
OTX2
GSX2
DLX5
ARX
DLX2
NKX2.1
NR2F1
PBX1
DLX1
LHX6
SP9
percent peaks in PSC
distal
proximal
f
LHX6
ARX
DLX5
DLX1
GSX2
SP9
DLX2
NKX2.1
OTX2
PBX1
NR2F1
ASCL1
percent peaks in loops
distal
proximal
g
number of occurrences
interaction ensemble size
h
occurrences with TFs bound
100%
90%
80%
70%
60%
50%
interaction ensemble size
i
chromatin state
outside
inside
PSC
j
chromatin state
Genome %
CpG Island
Exon
Gene
TES
TSS
TSS 2kb
H3K27me3
H3K4me3
H3K27ac
H3K4me1
EnhLo
EnhTx
Enh
EnhPoi
EnhBiv
ReprPC
Quies
TxWk
TssBiv
