## Supplementary Fig 2 for "Combinatorial transcription factor binding encodes cis-regulatory wiring of forebrain GABAergic neurogenesis"

### Supplementary Figure S2

**Combinatorial TF binding assignment for pREs at the *Arx* locus and primary TF motif by cluster. (a)** *Arx* locus on the X chromosome showing TF ChIP-seq signal and cluster assignment.  Region shows intermixing of pREs with different combinatorial binding patterns, as well as enrichment of broadly-bound pREs. Also shown are six regions previous tested in the VISTA enhancer database (https://enhancer.lbl.gov). 4/6 tested regions showed enhancer activity in E11.5 mouse forebrain. **(b)** Average motif positional coverage plots 300 bp in each direction from locus center for core TF motifs by pRE cluster. Scaled by row for comparable relative motif positional distribution within clusters. Comparison of positional motif enrichment (here) with proportional motif occurrence (Figure 2f) enables discernment of strong centered primary binding (motif presence with central positional tendency), dispersed but still putative primary DNA binding (motif presence, no central positional tendency), or indirect interaction (no motif presence, no central positional tendency).

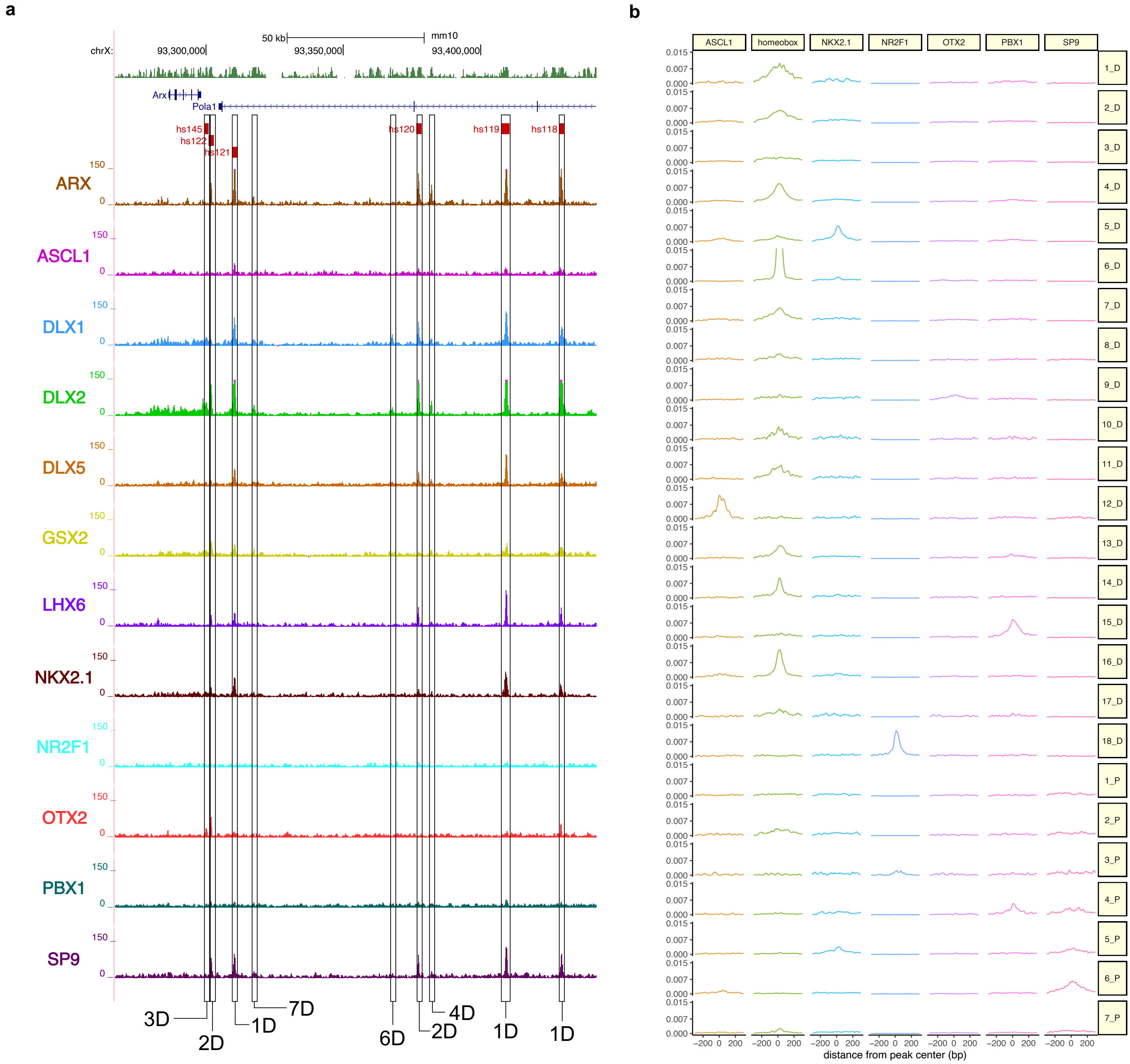

a
b
50 kb
mm10
chrX:
93,300,000
93,350,000
93,400,000
Arx
Pola1
hs145
hs122
hs121
hs120
hs119
hs118
ARX
ASCL1
DLX1
DLX2
DLX5
GSX2
LHX6
NKX2.1
NR2F1
OTX2
PBX1
SP9
150
0
3D
2D
1D
7D
6D
2D
4D
1D
1D
ASCL1
homeobox
NKX2.1
NR2F1
OTX2
PBX1
SP9
0.015
0.007
0.000
1_D
2_D
3_D
4_D
5_D
6_D
7_D
8_D
9_D
10_D
11_D
12_D
13_D
14_D
15_D
16_D
17_D
18_D
1_P
2_P
3_P
4_P
5_P
6_P
7_P
−200 0 200
distance from peak center (bp)
