## Supplementary Fig 3 for "Combinatorial transcription factor binding encodes cis-regulatory wiring of forebrain GABAergic neurogenesis"

### Supplementary Figure S3

**Additional features of combinatorial pRE clusters. (a)** Heatmap showing select Gene Ontology terms associated with genes with promoters are localized in the chromatin interaction ensembles for pRE clusters (adjusted p-value and q-value < 0.05). Scale is relative enrichment compared to expected. **(b)** Heatmap showing relative motif enrichment across clusters for select pRE cluster enriched HOMER TF motifs (Foxo1, TATA box, Oct4, Sox2, Olig2, E2A, NFY, and KLF14). Scale is proportion of peaks with motif. **(c)** Heatmap showing representation of previously identified activating/repressive regulatory elements (a.RE/r.RE) from knockout mouse studies of DLX1/2 and NKX2.1 TFs. Plot shows the enrichment of knockout sensitive a.RE/r.RE loci across clusters, when compared with expected by chance (natural log scale). Red color denotes overrepresentation, whereas blue is associated with underrepresentation; gray denotes too low number to assess. Color intensity is associated with extent of (over/under)representation. See main text for further details. **(d)** Boxplot showing relative likelihood of pREs within ensembles for DLX and NKX2.1, showing that DLX a.REs are twice as likely associated with the ensembles than not. Conversely, DLX r.REs are twice as likely not to be present in the ensembles. For NKX2.1, both a.REs and r.REs are more likely to be associated with ensembles. **(e-g)** Comparison of BG TF-bound pREs with TF-bound pREs previously reported in a study that characterized binding of five TFs (EMX2, LHX2, NR2F1, PAX6, and PBX1) in relatively stage-matched developing cortex/pallium[1]. (e) Pie chart showing the overlap of TF-bound loci in GEs with TF-bound loci in embryonic cortex. (f) Bar plot depicting the percent of cortex TF ChIP-seq peaks overlapping with GEs across clusters. (g) Bar plot depicting the proportion of BG pREs by cluster that overlapped with cortex TF ChIP-seq peaks. The highest proportion overlap among BG pRE clusters was for 1_D. Note that 3_P and 18_D are NR2F1-specific BG pRE clusters, and overlap captures strong overlap in binding targets for NR2F1 between developing BG and cortex.


1. Ypsilanti, A. R. *et al.* Transcriptional network orchestrating regional patterning of cortical progenitors. *Proc. Natl. Acad. Sci.* **118**, e2024795118 (2021).

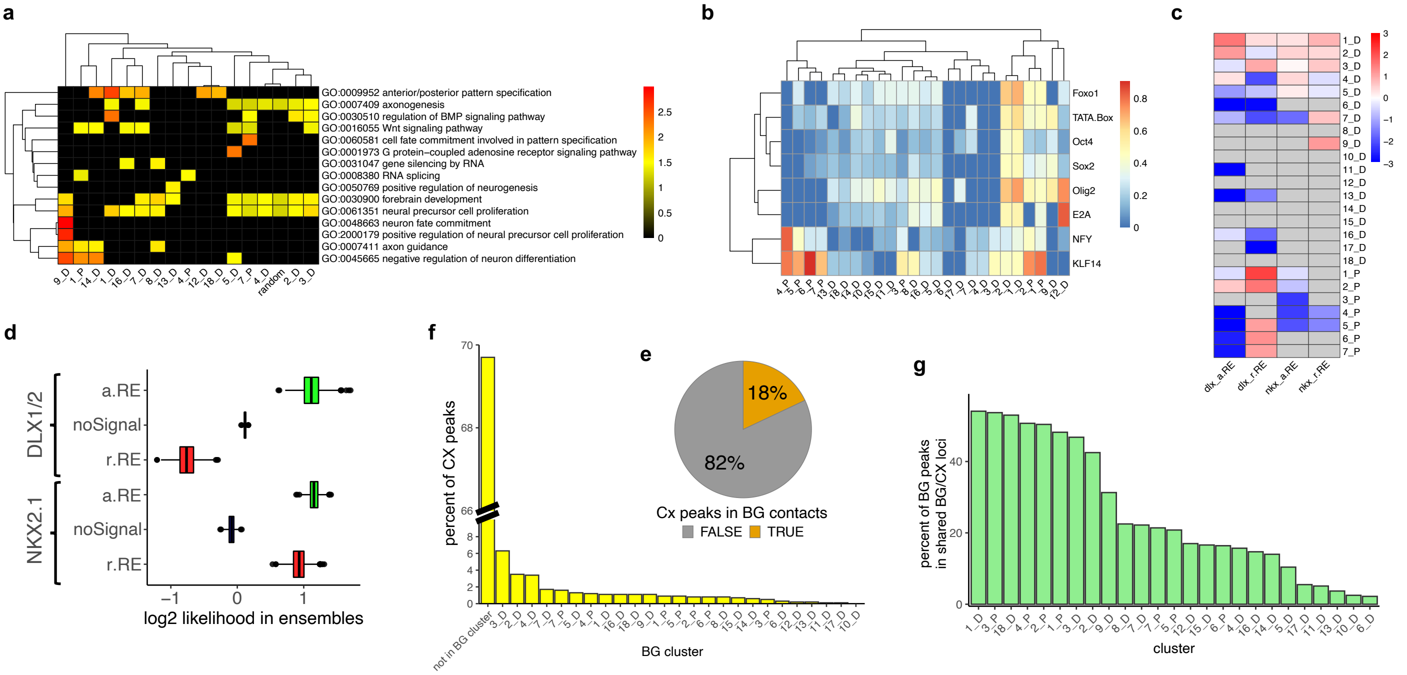

a
GO:0009952 anterior/posterior pattern specification
GO:0007409 axonogenesis
GO:0030510 regulation of BMP signaling pathway
GO:0016055 Wnt signaling pathway
GO:0060581 cell fate commitment involved in pattern specification
GO:0001973 G protein−coupled adenosine receptor signaling pathway
GO:0031047 gene silencing by RNA
GO:0008380 RNA splicing
GO:0050769 positive regulation of neurogenesis
GO:0030900 forebrain development
GO:0061351 neural precursor cell proliferation
GO:0048663 neuron fate commitment
GO:2000179 positive regulation of neural precursor cell proliferation
GO:0007411 axon guidance
GO:0045665 negative regulation of neuron differentiation
b
Foxo1
TATA.Box
Oct4
Sox2
Olig2
E2A
NFY
KLF14
c
dlx_a.RE
dlx_r.RE
nkx_a.RE
nkx_r.RE
d
DLX1/2
NKX2.1
a.RE
noSignal
r.RE
log2 likelihood in ensembles
e
18%
82%
Cx peaks in BG contacts
FALSE
TRUE
f
percent of CX peaks
not in BG cluster
BG cluster
g
percent of BG peaks in shared BG/CX loci
cluster
