## Supplementary Fig 4 for "Combinatorial transcription factor binding encodes cis-regulatory wiring of forebrain GABAergic neurogenesis"

### Supplementary Figure S4

**Summary of results for 84 pREs characterized here via transgenic mouse assays.** VISTA enhancer name (*mmXXXX*) is written in light blue to the left, below is internal enhancer name (*mJRXX*) as well as age of the stained and sectioned embryos. The success ratio is listed in dark blue to the right of the name (6/7 depicts 6 embryos with forebrain activity out of 7 embryos tested in total). Green and red bars show the normalized ChIP-seq signal of TFs in either BG (green, E13.5) or from cortex (CX, red, E12.5) TF ChIP-seq[1]. Darker color represents a stronger ChIP-Seq peak. The specific TF used in the ChIP-Seq experiments are shown at the top. Below each bar is a schema showing the tested enhancer location in light blue relative to the regulated gene, generally defined via H3K4me3 PLAC-seq interaction. The grey arrow depicts the orientation of target gene transcription. To the right of the gene schema is a wholemount (WM) of a representative of the *LacZ* expression, next to the WM are three sections of the specific WM shown. Only the WM are shown for pREs not having clearly reproducible *LacZ* activity in the forebrain. The pREs are listed based on their cluster number (bold numbers to the left, i.e., 1_D first, 2_D second, etc.) and within each cluster number the enhancers are sorted by activity: exclusively subpallium first (light green bar to the right), subpallium+pallium second (purple bar to the right), and exclusively pallium last (dark pink bar to the right). pREs marked with an * were not included in further studies due to a positive rate below 30%. *mm1199*, *mm1276*, *mm1422*, *mm1423*, *mm1437*, *mm1787*, *mm1788*, *mm1804*, *mm1814*, *mm1816*, *mm1819* are published in [1]; *mm1196* is published in [2]; *mm1203* is published in [3]; *mm1428* and *mm1429* are published in [4]. Full transgenic enhancer assay results available at https://enhancer.lbl.gov.


1. Ypsilanti, A. R. *et al.* Transcriptional network orchestrating regional patterning of cortical progenitors. *Proc. Natl. Acad. Sci.* **118**, e2024795118 (2021).
2. Pla, R. *et al.* Dlx1 and Dlx2 Promote Interneuron GABA Synthesis, Synaptogenesis, and Dendritogenesis. *Cereb. Cortex* **28**, 3797–3815 (2018).
3. Lindtner, S. *et al.* Genomic Resolution of DLX-Orchestrated Transcriptional Circuits Driving Development of Forebrain GABAergic Neurons. *Cell Rep.* **28**, 2048-2063.e8 (2019).
4. Sandberg, M. *et al.* Transcriptional Networks Controlled by NKX2-1 in the Development of Forebrain GABAergic Neurons. *Neuron* **91**, 1260–1275 (2016).

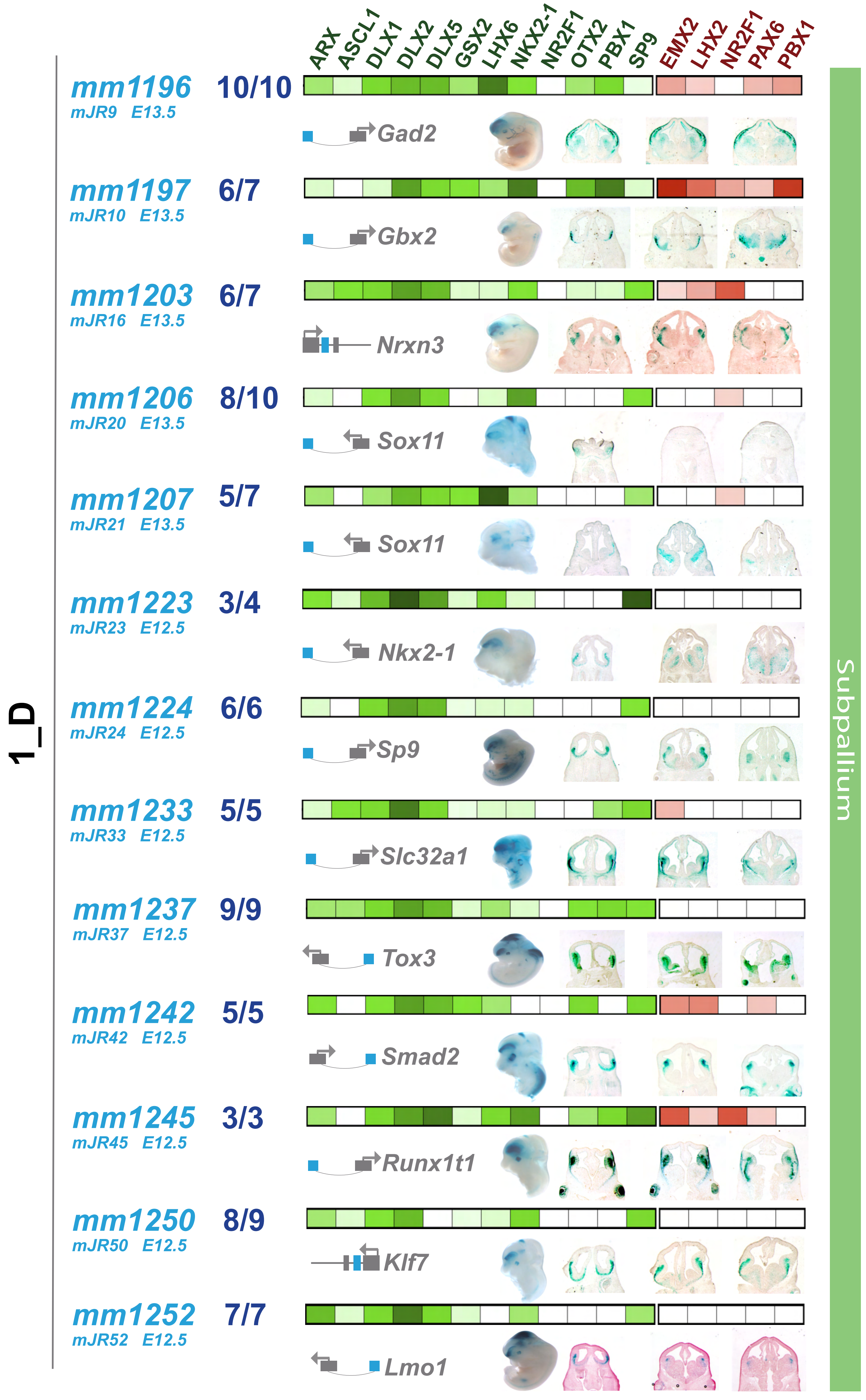

1_D
ARX ASCL1 DLX1 DLX2 DLX5 GSX2 LHX6 NKX2-1 NR2F1 OTX2 PBX1 SP9
EMX2 LHX2 NR2F1 PAX6 PBX1
Subpallium
mm1196 mJR9 E13.5 10/10 Gad2
mm1197 mJR10 E13.5 6/7 Gbx2
mm1203 mJR16 E13.5 6/7 Nrxn3
mm1206 mJR20 E13.5 8/10 Sox11
mm1207 mJR21 E13.5 5/7 Sox11
mm1223 mJR23 E12.5 3/4 Nkx2-1
mm1224 mJR24 E12.5 6/6 Sp9
mm1233 mJR33 E12.5 5/5 Slc32a1
mm1237 mJR37 E12.5 9/9 Tox3
mm1242 mJR42 E12.5 5/5 Smad2
mm1245 mJR45 E12.5 3/3 Runx1t1
mm1250 mJR50 E12.5 8/9 Klf7
mm1252 mJR52 E12.5 7/7 Lmo1

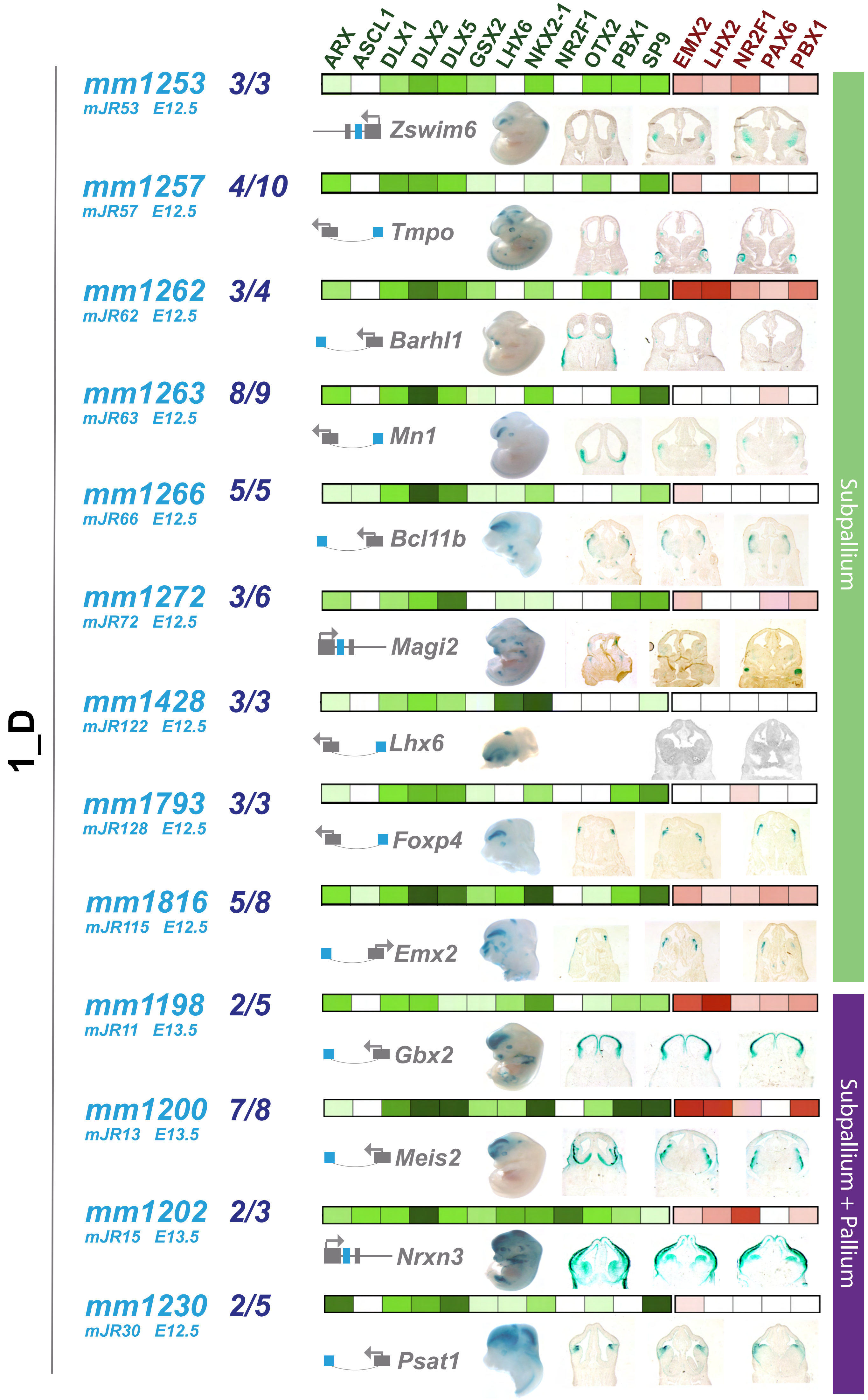
1_D
ARX ASCL1 DLX1 DLX2 DLX5 GSX2 LHX6 NKX2-1 NR2F1 OTX2 PBX1 SP9
EMX2 LHX2 NR2F1 PAX6 PBX1
mm1253 3/3
mJR53 E12.5
Zswim6
mm1257 4/10
mJR57 E12.5
Tmpo
mm1262 3/4
mJR62 E12.5
Barhl1
mm1263 8/9
mJR63 E12.5
Mn1
mm1266 5/5
mJR66 E12.5
Bcl11b
mm1272 3/6
mJR72 E12.5
Magi2
mm1428 3/3
mJR122 E12.5
Lhx6
mm1793 3/3
mJR128 E12.5
Foxp4
mm1816 5/8
mJR115 E12.5
Emx2
mm1198 2/5
mJR11 E13.5
Gbx2
mm1200 7/8
mJR13 E13.5
Meis2
mm1202 2/3
mJR15 E13.5
Nrxn3
mm1230 2/5
mJR30 E12.5
Psat1
Subpallium
Subpallium + Pallium

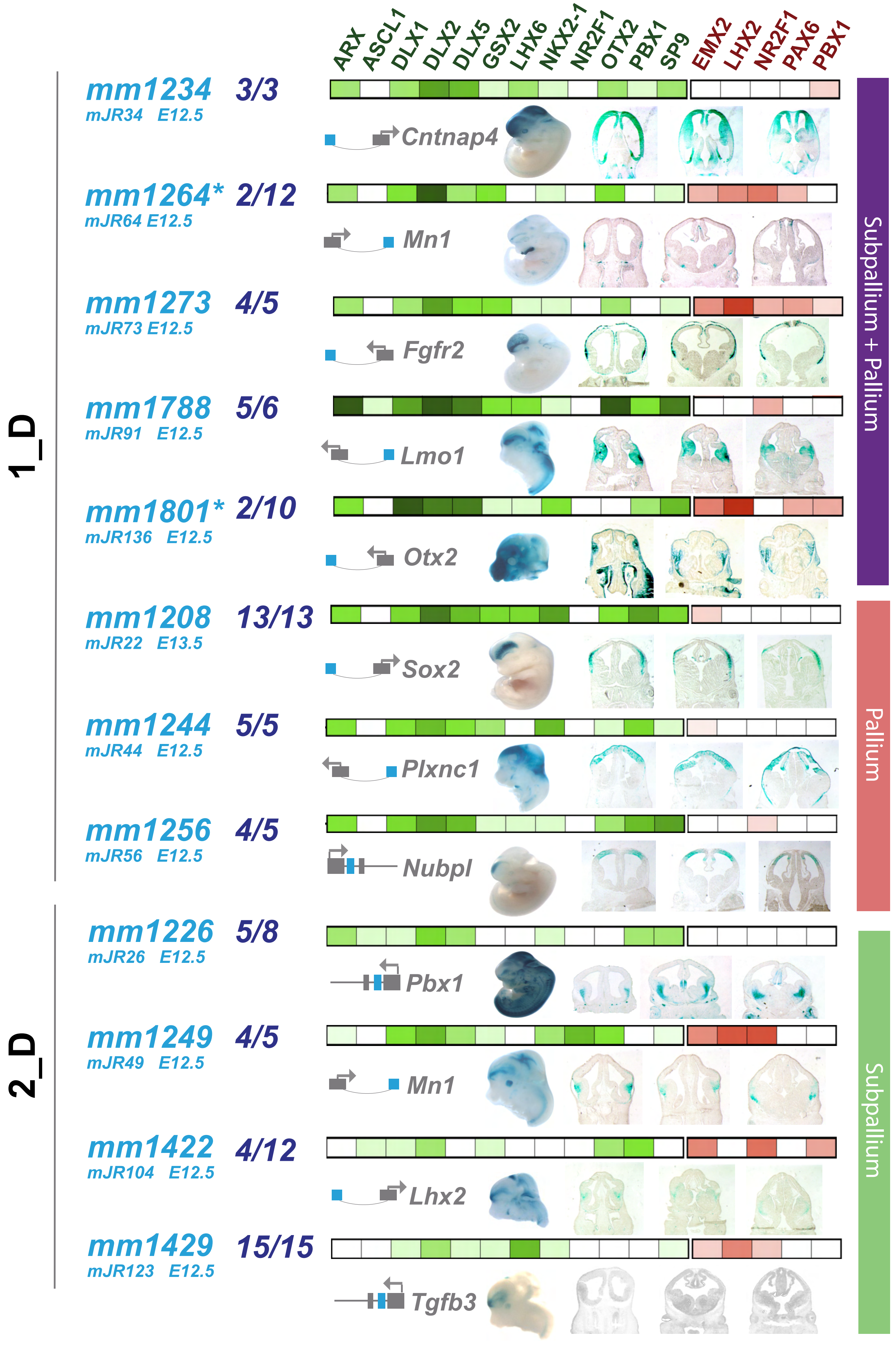

ARX
ASCL1
DLX1
DLX2
DLX5
GSX2
LHX6
NKX2-1
NR2F1
OTX2
PBX1
SP9
EMX2
LHX2
NR2F1
PAX6
PBX1
1_D
2_D
mm1234 3/3
mJR34 E12.5
Cntnap4
mm1264* 2/12
mJR64 E12.5
Mn1
mm1273 4/5
mJR73 E12.5
Fgfr2
mm1788 5/6
mJR91 E12.5
Lmo1
mm1801* 2/10
mJR136 E12.5
Otx2
mm1208 13/13
mJR22 E13.5
Sox2
mm1244 5/5
mJR44 E12.5
Plxnc1
mm1256 4/5
mJR56 E12.5
Nubpl
mm1226 5/8
mJR26 E12.5
Pbx1
mm1249 4/5
mJR49 E12.5
Mn1
mm1422 4/12
mJR104 E12.5
Lhx2
mm1429 15/15
mJR123 E12.5
Tgfb3
Subpallium + Pallium
Pallium
Subpallium

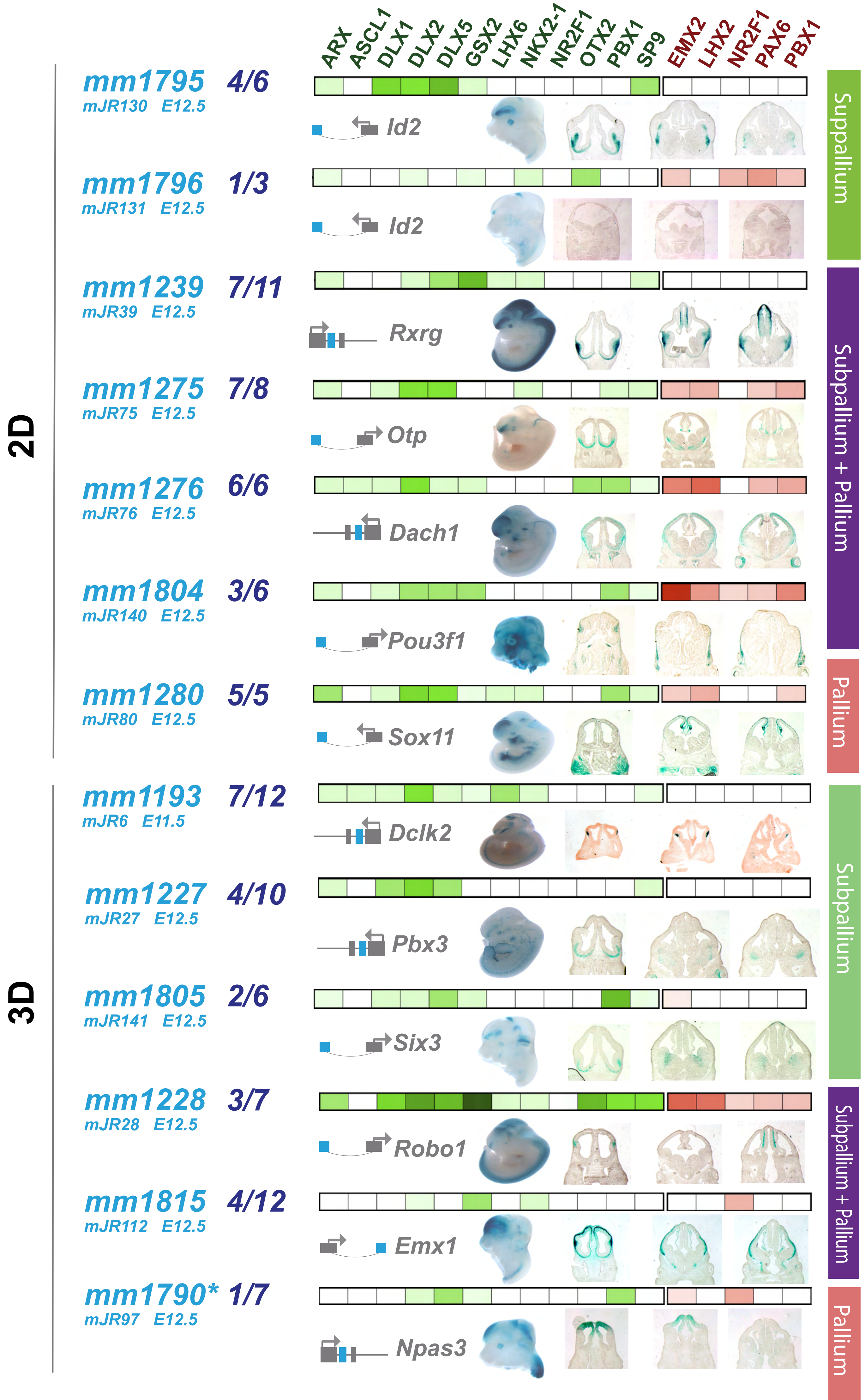
ARX ASCL1 DLX1 DLX2 DLX5 GSX2 LHX6 NKX2-1 NR2F1 OTX2 PBX1 SP9 EMX2 LHX2 NR2F1 PAX6 PBX1
2D
3D
mm1795 4/6
mJR130 E12.5
Id2
mm1796 1/3
mJR131 E12.5
Id2
mm1239 7/11
mJR39 E12.5
Rxrg
mm1275 7/8
mJR75 E12.5
Otp
mm1276 6/6
mJR76 E12.5
Dach1
mm1804 3/6
mJR140 E12.5
Pou3f1
mm1280 5/5
mJR80 E12.5
Sox11
mm1193 7/12
mJR6 E11.5
Dclk2
mm1227 4/10
mJR27 E12.5
Pbx3
mm1805 2/6
mJR141 E12.5
Six3
mm1228 3/7
mJR28 E12.5
Robo1
mm1815 4/12
mJR112 E12.5
Emx1
mm1790* 1/7
mJR97 E12.5
Npas3
Subpallium
Subpallium + Pallium
Pallium
Subpallium
Subpallium + Pallium
Pallium

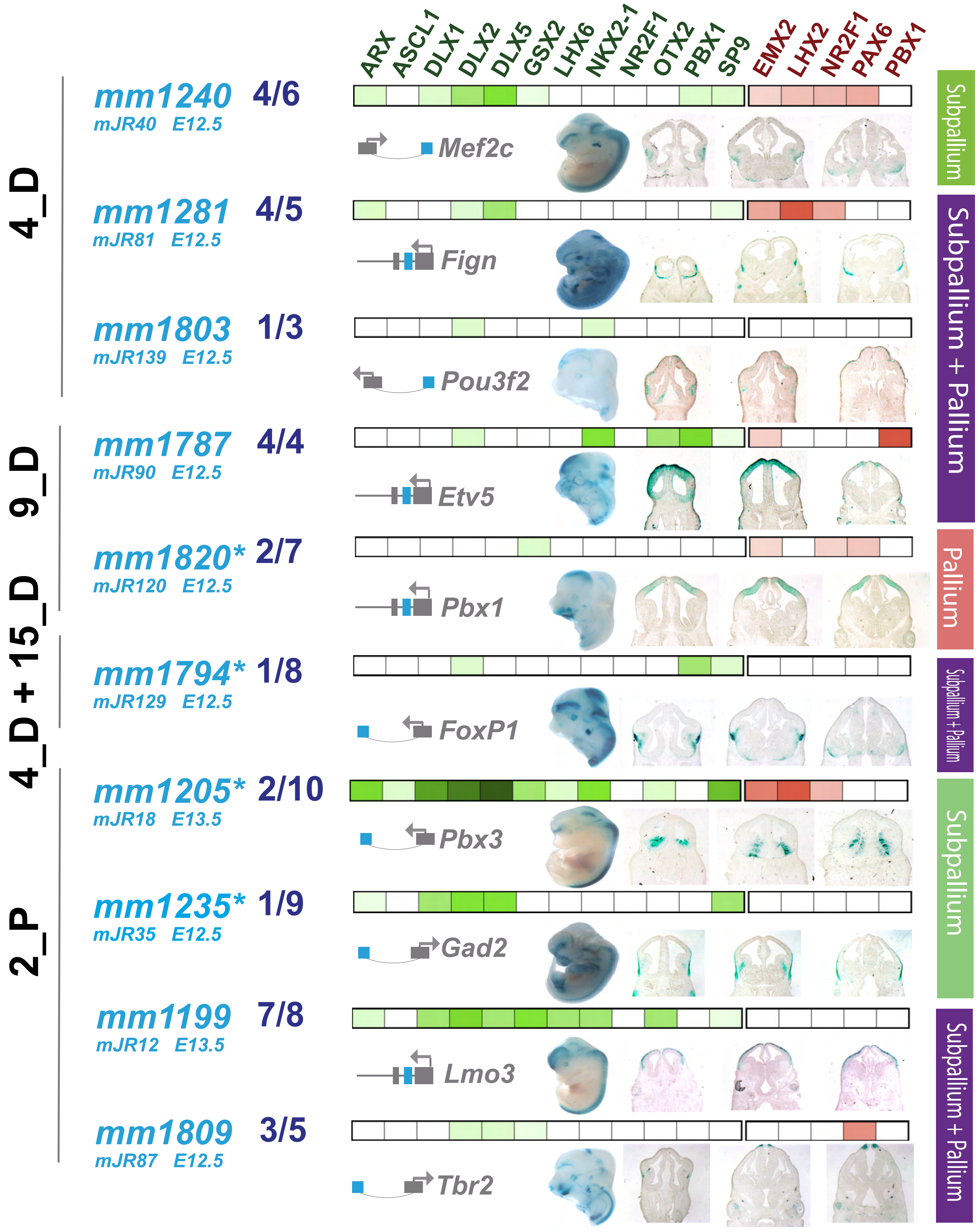

ARX
ASCL1
DLX1
DLX2
DLX5
GSX2
LHX6
NKX2-1
NR2F1
OTX2
PBX1
SP9
EMX2
LHX2
NR2F1
PAX6
PBX1
4_D
9_D
4_D + 15_D
2_P
mm1240 4/6
mJR40 E12.5
Mef2c
mm1281 4/5
mJR81 E12.5
Fign
mm1803 1/3
mJR139 E12.5
Pou3f2
mm1787 4/4
mJR90 E12.5
Etv5
mm1820* 2/7
mJR120 E12.5
Pbx1
mm1794* 1/8
mJR129 E12.5
FoxP1
mm1205* 2/10
mJR18 E13.5
Pbx3
mm1235* 1/9
mJR35 E12.5
Gad2
mm1199 7/8
mJR12 E13.5
Lmo3
mm1809 3/5
mJR87 E12.5
Tbr2
Subpallium
Subpallium + Pallium
Pallium
Subpallium + Pallium
Subpallium
Subpallium + Pallium

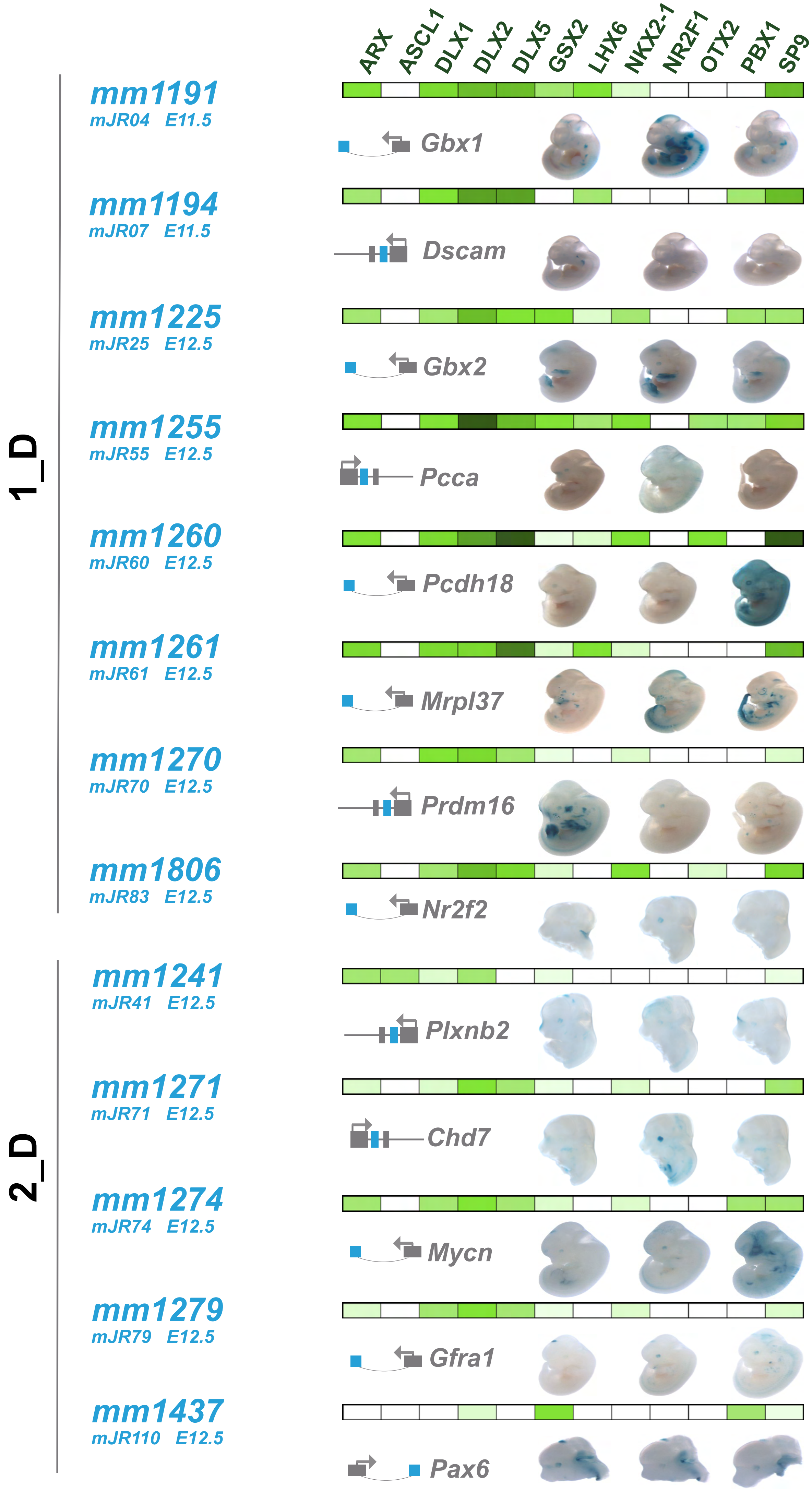
ARX
ASCL1
DLX1
DLX2
DLX5
GSX2
LHX6
NKX2-1
NR2F1
OTX2
PBX1
SP9
1_D
mm1191
mJR04 E11.5
Gbx1
mm1194
mJR07 E11.5
Dscam
mm1225
mJR25 E12.5
Gbx2
mm1255
mJR55 E12.5
Pcca
mm1260
mJR60 E12.5
Pcdh18
mm1261
mJR61 E12.5
Mrpl37
mm1270
mJR70 E12.5
Prdm16
mm1806
mJR83 E12.5
Nr2f2
2_D
mm1241
mJR41 E12.5
Plxnb2
mm1271
mJR71 E12.5
Chd7
mm1274
mJR74 E12.5
Mycn
mm1279
mJR79 E12.5
Gfra1
mm1437
mJR110 E12.5
Pax6

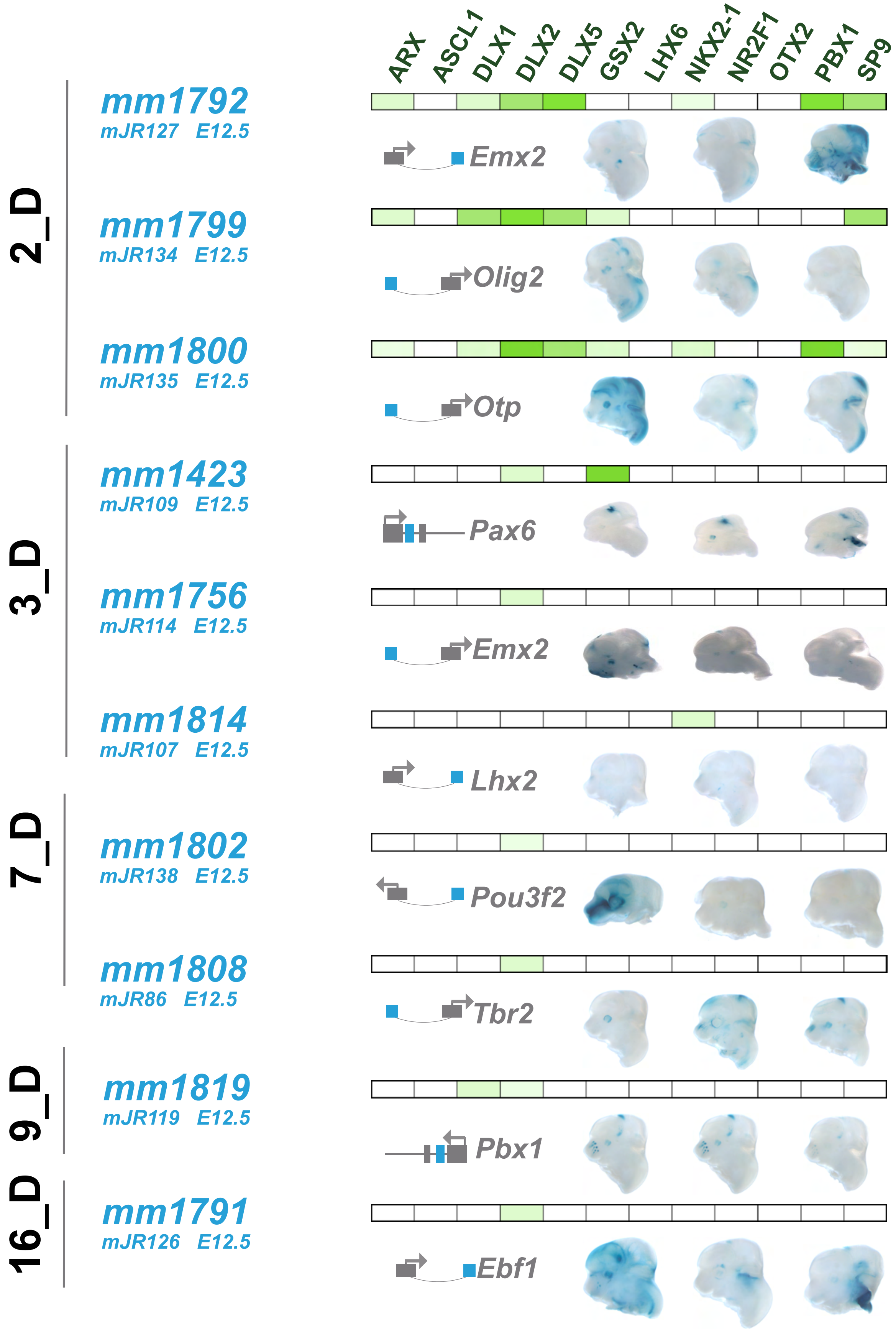
ARX
ASCL1
DLX1
DLX2
DLX5
GSX2
LHX6
NKX2-1
NR2F1
OTX2
PBX1
SP9
2_D
mm1792
mJR127 E12.5
Emx2
mm1799
mJR134 E12.5
Olig2
mm1800
mJR135 E12.5
Otp
3_D
mm1423
mJR109 E12.5
Pax6
mm1756
mJR114 E12.5
Emx2
mm1814
mJR107 E12.5
Lhx2
7_D
mm1802
mJR138 E12.5
Pou3f2
mm1808
mJR86 E12.5
Tbr2
9_D
mm1819
mJR119 E12.5
Pbx1
16_D
mm1791
mJR126 E12.5
Ebf1
