## Supplementary Fig 5 for "Combinatorial transcription factor binding encodes cis-regulatory wiring of forebrain GABAergic neurogenesis"

### Supplementary Figure S5

**Average base-level PhyloP vertebrate evolutionary conservation of motifs within pRE clusters. (a)** Average base-level conservation for Primary TF motifs identified in this study for each pRE cluster. Values calculated by centering on each motif instance for all loci within pRE cluster and then calculating average value at that relative position. Left plots show row centered and scaled normalized Z-score representation of PhyloP scores. Right plots show raw average PhyloP values. Relative plots useful for comparing change in conservation between motif and neighboring sequence, whereas raw values capture background conservation versus change within motif. **(b)** Base-level conservation across all HOMER motifs that are enriched within 1_D cluster. Calculated in same manner as for (a).

a

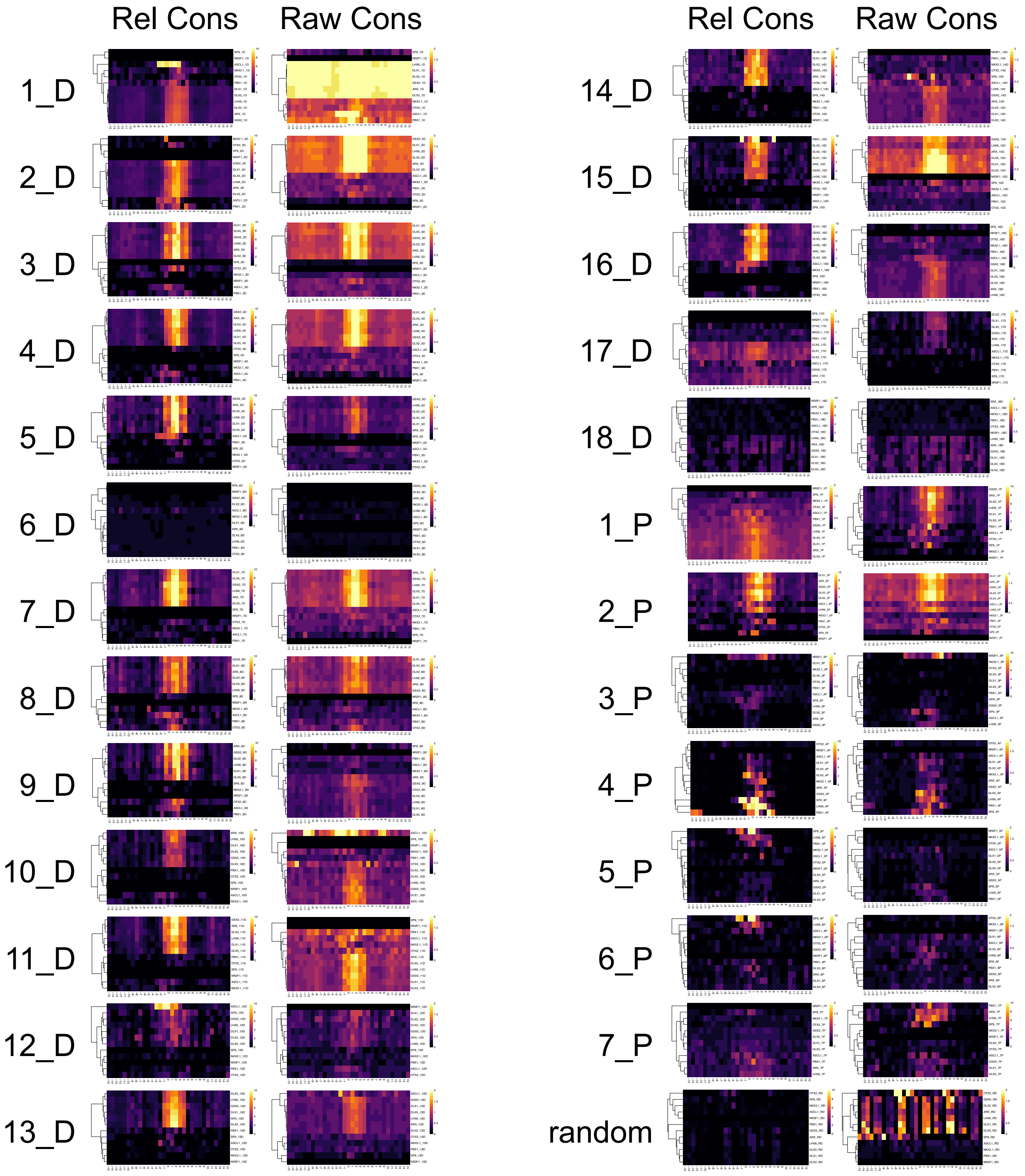

Rel Cons
Raw Cons
Rel Cons
Raw Cons
1_D
2_D
3_D
4_D
5_D
6_D
7_D
8_D
9_D
10_D
11_D
12_D
13_D
14_D
15_D
16_D
17_D
18_D
1_P
2_P
3_P
4_P
5_P
6_P
7_P
random

**b**

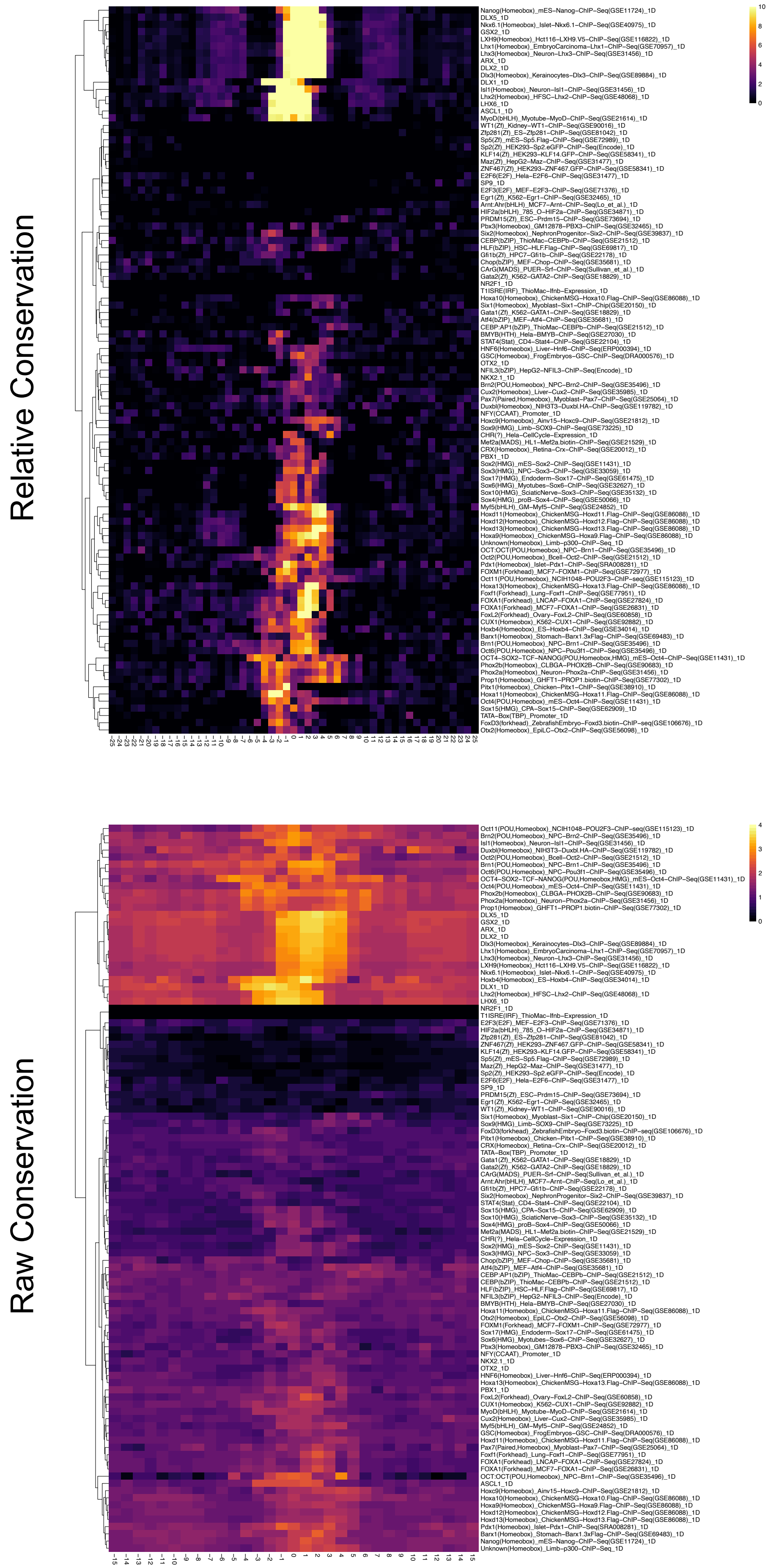
Relative Conservation
Nanog(Homeobox)_mES-Nanog-ChIP-Seq(GSE11724)_1D
DLX5_1D
Nkx6.1(Homeobox)_Islet-Nkx6.1-ChIP-Seq(GSE40975)_1D
GSX2_1D
LXH9(Homeobox)_Hct116-LXH9.V5-ChIP-Seq(GSE116822)_1D
Lhx1(Homeobox)_EmbryoCarcinoma-Lhx1-ChIP-Seq(GSE70957)_1D
Lhx3(Homeobox)_Neuron-Lhx3-ChIP-Seq(GSE31456)_1D
ARX_1D
DLX2_1D
Dlx3(Homeobox)_Kerainocytes-Dlx3-ChIP-Seq(GSE89884)_1D
DLX1_1D
Isl1(Homeobox)_Neuron-Isl1-ChIP-Seq(GSE31456)_1D
Lhx2(Homeobox)_HFSC-Lhx2-ChIP-Seq(GSE48068)_1D
LHX6_1D
ASCL1_1D
MyoD(bHLH)_Myotube-MyoD-ChIP-Seq(GSE21614)_1D
WT1(Zf)_Kidney-WT1-ChIP-Seq(GSE90016)_1D
Zfp281(Zf)_ES-Zfp281-ChIP-Seq(GSE81042)_1D
Sp5(Zf)_mES-Sp5.Flag-ChIP-Seq(GSE72989)_1D
Sp2(Zf)_HEK293-Sp2.eGFP-ChIP-Seq(Encode)_1D
KLF14(Zf)_HEK293-KLF14.GFP-ChIP-Seq(GSE58341)_1D
Maz(Zf)_HepG2-Maz-ChIP-Seq(GSE31477)_1D
ZNF467(Zf)_HEK293-ZNF467.GFP-ChIP-Seq(GSE58341)_1D
E2F6(E2F)_Hela-E2F6-ChIP-Seq(GSE31477)_1D
SP9_1D
E2F3(E2F)_MEF-E2F3-ChIP-Seq(GSE71376)_1D
Egr1(Zf)_K562-Egr1-ChIP-Seq(GSE32465)_1D
Arnt:Ahr(bHLH)_MCF7-Arnt-ChIP-Seq(Lo_et_al.)_1D
HIF2a(bHLH)_785_O-HIF2a-ChIP-Seq(GSE34871)_1D
PRDM15(Zf)_ESC-Prdm15-ChIP-Seq(GSE73694)_1D
Pbx3(Homeobox)_GM12878-PBX3-ChIP-Seq(GSE32465)_1D
Six2(Homeobox)_NephronProgenitor-Six2-ChIP-Seq(GSE39837)_1D
CEBP(bZIP)_ThioMac-CEBPb-ChIP-Seq(GSE21512)_1D
HLF(bZIP)_HSC-HLF.Flag-ChIP-Seq(GSE69817)_1D
Gfi1b(Zf)_HPC7-Gfi1b-ChIP-Seq(GSE22178)_1D
Chop(bZIP)_MEF-Chop-ChIP-Seq(GSE35681)_1D
CArG(MADS)_PUER-Srf-ChIP-Seq(Sullivan_et_al.)_1D
Gata2(Zf)_K562-GATA2-ChIP-Seq(GSE18829)_1D
NR2F1_1D
T1ISRE(IRF)_ThioMac-Ifnb-Expression_1D
Hoxa10(Homeobox)_ChickenMSG-Hoxa10.Flag-ChIP-Seq(GSE86088)_1D
Six1(Homeobox)_Myoblast-Six1-ChIP-Chip(GSE20150)_1D
Gata1(Zf)_K562-GATA1-ChIP-Seq(GSE18829)_1D
Atf4(bZIP)_MEF-Atf4-ChIP-Seq(GSE35681)_1D
CEBP:AP1(bZIP)_ThioMac-CEBPb-ChIP-Seq(GSE21512)_1D
BMYB(HTH)_Hela-BMYB-ChIP-Seq(GSE27030)_1D
STAT4(Stat)_CD4-Stat4-ChIP-Seq(GSE22104)_1D
HNF6(Homeobox)_Liver-Hnf6-ChIP-Seq(ERP000394)_1D
GSC(Homeobox)_FrogEmbryos-GSC-ChIP-Seq(DRA000576)_1D
OTX2_1D
NFIL3(bZIP)_HepG2-NFIL3-ChIP-Seq(Encode)_1D
NKX2.1_1D
Brn2(POU,Homeobox)_NPC-Brn2-ChIP-Seq(GSE35496)_1D
Cux2(Homeobox)_Liver-Cux2-ChIP-Seq(GSE35985)_1D
Pax7(Paired,Homeobox)_Myoblast-Pax7-ChIP-Seq(GSE25064)_1D
Duxbl(Homeobox)_NIH3T3-Duxbl.HA-ChIP-Seq(GSE119782)_1D
NFY(CCAAT)_Promoter_1D
Hoxc9(Homeobox)_Ainv15-Hoxc9-ChIP-Seq(GSE21812)_1D
Sox9(HMG)_Limb-SOX9-ChIP-Seq(GSE73225)_1D
CHR(?)_Hela-CellCycle-Expression_1D
Mef2a(MADS)_HL1-Mef2a.biotin-ChIP-Seq(GSE21529)_1D
CRX(Homeobox)_Retina-Crx-ChIP-Seq(GSE20012)_1D
PBX1_1D
Sox2(HMG)_mES-Sox2-ChIP-Seq(GSE11431)_1D
Sox3(HMG)_NPC-Sox3-ChIP-Seq(GSE33059)_1D
Sox17(HMG)_Endoderm-Sox17-ChIP-Seq(GSE61475)_1D
Sox6(HMG)_Myotubes-Sox6-ChIP-Seq(GSE32627)_1D
Sox10(HMG)_SciaticNerve-Sox3-ChIP-Seq(GSE35132)_1D
Sox4(HMG)_proB-Sox4-ChIP-Seq(GSE50066)_1D
Myf5(bHLH)_GM-Myf5-ChIP-Seq(GSE24852)_1D
Hoxd11(Homeobox)_ChickenMSG-Hoxd11.Flag-ChIP-Seq(GSE86088)_1D
Hoxd12(Homeobox)_ChickenMSG-Hoxd12.Flag-ChIP-Seq(GSE86088)_1D
Hoxd13(Homeobox)_ChickenMSG-Hoxd13.Flag-ChIP-Seq(GSE86088)_1D
Hoxa9(Homeobox)_ChickenMSG-Hoxa9.Flag-ChIP-Seq(GSE86088)_1D
Unknown(Homeobox)_Limb-p300-ChIP-Seq_1D
OCT:OCT(POU,Homeobox)_NPC-Brn1-ChIP-Seq(GSE35496)_1D
Oct2(POU,Homeobox)_Bcell-Oct2-ChIP-Seq(GSE21512)_1D
Pdx1(Homeobox)_Islet-Pdx1-ChIP-Seq(SRA008281)_1D
FOXM1(Forkhead)_MCF7-FOXM1-ChIP-Seq(GSE72977)_1D
Oct11(POU,Homeobox)_NCIH1048-POU2F3-ChIP-seq(GSE115123)_1D
Hoxa13(Homeobox)_ChickenMSG-Hoxa13.Flag-ChIP-Seq(GSE86088)_1D
Foxf1(Forkhead)_Lung-Foxf1-ChIP-Seq(GSE77951)_1D
FOXA1(Forkhead)_LNCAP-FOXA1-ChIP-Seq(GSE27824)_1D
FOXA1(Forkhead)_MCF7-FOXA1-ChIP-Seq(GSE26831)_1D
FoxL2(Forkhead)_Ovary-FoxL2-ChIP-Seq(GSE60858)_1D
CUX1(Homeobox)_K562-CUX1-ChIP-Seq(GSE92882)_1D
Hoxb4(Homeobox)_ES-Hoxb4-ChIP-Seq(GSE34014)_1D
Barx1(Homeobox)_Stomach-Barx1.3xFlag-ChIP-Seq(GSE69483)_1D
Brn1(POU,Homeobox)_NPC-Brn1-ChIP-Seq(GSE35496)_1D
Oct6(POU,Homeobox)_NPC-Pou3f1-ChIP-Seq(GSE35496)_1D
OCT4-SOX2-TCF-NANOG(POU,Homeobox,HMG)_mES-Oct4-ChIP-Seq(GSE11431)_1D
Phox2b(Homeobox)_CLBGA-PHOX2B-ChIP-Seq(GSE90683)_1D
Phox2a(Homeobox)_Neuron-Phox2a-ChIP-Seq(GSE31456)_1D
Prop1(Homeobox)_GHFT1-PROP1.biotin-ChIP-Seq(GSE77302)_1D
Pitx1(Homeobox)_Chicken-Pitx1-ChIP-Seq(GSE38910)_1D
Hoxa11(Homeobox)_ChickenMSG-Hoxa11.Flag-ChIP-Seq(GSE86088)_1D
Oct4(POU,Homeobox)_mES-Oct4-ChIP-Seq(GSE11431)_1D
Sox15(HMG)_CPA-Sox15-ChIP-Seq(GSE62909)_1D
TATA-Box(TBP)_Promoter_1D
FoxD3(forkhead)_ZebrafishEmbryo-Foxd3.biotin-ChIP-seq(GSE106676)_1D
Otx2(Homeobox)_EpiLC-Otx2-ChIP-Seq(GSE56098)_1D
-25 -24 -23 -22 -21 -20 -19 -18 -17 -16 -15 -14 -13 -12 -11 -10 -9 -8 -7 -6 -5 -4 -3 -2 -1 0 1 2 3 4 5 6 7 8 9 10 11 12 13 14 15 16 17 18 19 20 21 22 23 24 25
10 8 6 4 2 0
Raw Conservation
Oct11(POU,Homeobox)_NCIH1048-POU2F3-ChIP-seq(GSE115123)_1D
Brn2(POU,Homeobox)_NPC-Brn2-ChIP-Seq(GSE35496)_1D
Isl1(Homeobox)_Neuron-Isl1-ChIP-Seq(GSE31456)_1D
Duxbl(Homeobox)_NIH3T3-Duxbl.HA-ChIP-Seq(GSE119782)_1D
Oct2(POU,Homeobox)_Bcell-Oct2-ChIP-Seq(GSE21512)_1D
Brn1(POU,Homeobox)_NPC-Brn1-ChIP-Seq(GSE35496)_1D
Oct6(POU,Homeobox)_NPC-Pou3f1-ChIP-Seq(GSE35496)_1D
OCT4-SOX2-TCF-NANOG(POU,Homeobox,HMG)_mES-Oct4-ChIP-Seq(GSE11431)_1D
Oct4(POU,Homeobox)_mES-Oct4-ChIP-Seq(GSE11431)_1D
Phox2b(Homeobox)_CLBGA-PHOX2B-ChIP-Seq(GSE90683)_1D
Phox2a(Homeobox)_Neuron-Phox2a-ChIP-Seq(GSE31456)_1D
Prop1(Homeobox)_GHFT1-PROP1.biotin-ChIP-Seq(GSE77302)_1D
DLX5_1D
GSX2_1D
ARX_1D
DLX2_1D
Dlx3(Homeobox)_Kerainocytes-Dlx3-ChIP-Seq(GSE89884)_1D
Lhx1(Homeobox)_EmbryoCarcinoma-Lhx1-ChIP-Seq(GSE70957)_1D
Lhx3(Homeobox)_Neuron-Lhx3-ChIP-Seq(GSE31456)_1D
LXH9(Homeobox)_Hct116-LXH9.V5-ChIP-Seq(GSE116822)_1D
Nkx6.1(Homeobox)_Islet-Nkx6.1-ChIP-Seq(GSE40975)_1D
Hoxb4(Homeobox)_ES-Hoxb4-ChIP-Seq(GSE34014)_1D
DLX1_1D
Lhx2(Homeobox)_HFSC-Lhx2-ChIP-Seq(GSE48068)_1D
LHX6_1D
NR2F1_1D
T1ISRE(IRF)_ThioMac-Ifnb-Expression_1D
E2F3(E2F)_MEF-E2F3-ChIP-Seq(GSE71376)_1D
HIF2a(bHLH)_785_O-HIF2a-ChIP-Seq(GSE34871)_1D
Zfp281(Zf)_ES-Zfp281-ChIP-Seq(GSE81042)_1D
ZNF467(Zf)_HEK293-ZNF467.GFP-ChIP-Seq(GSE58341)_1D
KLF14(Zf)_HEK293-KLF14.GFP-ChIP-Seq(GSE58341)_1D
Sp5(Zf)_mES-Sp5.Flag-ChIP-Seq(GSE72989)_1D
Maz(Zf)_HepG2-Maz-ChIP-Seq(GSE31477)_1D
Sp2(Zf)_HEK293-Sp2.eGFP-ChIP-Seq(Encode)_1D
E2F6(E2F)_Hela-E2F6-ChIP-Seq(GSE31477)_1D
SP9_1D
PRDM15(Zf)_ESC-Prdm15-ChIP-Seq(GSE73694)_1D
Egr1(Zf)_K562-Egr1-ChIP-Seq(GSE32465)_1D
WT1(Zf)_Kidney-WT1-ChIP-Seq(GSE90016)_1D
Six1(Homeobox)_Myoblast-Six1-ChIP-Chip(GSE20150)_1D
Sox9(HMG)_Limb-SOX9-ChIP-Seq(GSE73225)_1D
FoxD3(forkhead)_ZebrafishEmbryo-Foxd3.biotin-ChIP-seq(GSE106676)_1D
Pitx1(Homeobox)_Chicken-Pitx1-ChIP-Seq(GSE38910)_1D
CRX(Homeobox)_Retina-Crx-ChIP-Seq(GSE20012)_1D
TATA-Box(TBP)_Promoter_1D
Gata1(Zf)_K562-GATA1-ChIP-Seq(GSE18829)_1D
Gata2(Zf)_K562-GATA2-ChIP-Seq(GSE18829)_1D
CArG(MADS)_PUER-Srf-ChIP-Seq(Sullivan_et_al.)_1D
Arnt:Ahr(bHLH)_MCF7-Arnt-ChIP-Seq(Lo_et_al.)_1D
Gfi1b(Zf)_HPC7-Gfi1b-ChIP-Seq(GSE22178)_1D
Six2(Homeobox)_NephronProgenitor-Six2-ChIP-Seq(GSE39837)_1D
STAT4(Stat)_CD4-Stat4-ChIP-Seq(GSE22104)_1D
Sox15(HMG)_CPA-Sox15-ChIP-Seq(GSE62909)_1D
Sox10(HMG)_SciaticNerve-Sox3-ChIP-Seq(GSE35132)_1D
Sox4(HMG)_proB-Sox4-ChIP-Seq(GSE50066)_1D
Mef2a(MADS)_HL1-Mef2a.biotin-ChIP-Seq(GSE21529)_1D
CHR(?)_Hela-CellCycle-Expression_1D
Sox2(HMG)_mES-Sox2-ChIP-Seq(GSE11431)_1D
Sox3(HMG)_NPC-Sox3-ChIP-Seq(GSE33059)_1D
Chop(bZIP)_MEF-Chop-ChIP-Seq(GSE35681)_1D
Atf4(bZIP)_MEF-Atf4-ChIP-Seq(GSE35681)_1D
CEBP:AP1(bZIP)_ThioMac-CEBPb-ChIP-Seq(GSE21512)_1D
CEBP(bZIP)_ThioMac-CEBPb-ChIP-Seq(GSE21512)_1D
HLF(bZIP)_HSC-HLF.Flag-ChIP-Seq(GSE69817)_1D
NFIL3(bZIP)_HepG2-NFIL3-ChIP-Seq(Encode)_1D
BMYB(HTH)_Hela-BMYB-ChIP-Seq(GSE27030)_1D
Hoxa11(Homeobox)_ChickenMSG-Hoxa11.Flag-ChIP-Seq(GSE86088)_1D
Otx2(Homeobox)_EpiLC-Otx2-ChIP-Seq(GSE56098)_1D
FOXM1(Forkhead)_MCF7-FOXM1-ChIP-Seq(GSE72977)_1D
Sox17(HMG)_Endoderm-Sox17-ChIP-Seq(GSE61475)_1D
Sox6(HMG)_Myotubes-Sox6-ChIP-Seq(GSE32627)_1D
Pbx3(Homeobox)_GM12878-PBX3-ChIP-Seq(GSE32465)_1D
NFY(CCAAT)_Promoter_1D
NKX2.1_1D
OTX2_1D
HNF6(Homeobox)_Liver-Hnf6-ChIP-Seq(ERP000394)_1D
Hoxa13(Homeobox)_ChickenMSG-Hoxa13.Flag-ChIP-Seq(GSE86088)_1D
PBX1_1D
FoxL2(Forkhead)_Ovary-FoxL2-ChIP-Seq(GSE60858)_1D
CUX1(Homeobox)_K562-CUX1-ChIP-Seq(GSE92882)_1D
MyoD(bHLH)_Myotube-MyoD-ChIP-Seq(GSE21614)_1D
Cux2(Homeobox)_Liver-Cux2-ChIP-Seq(GSE35985)_1D
Myf5(bHLH)_GM-Myf5-ChIP-Seq(GSE24852)_1D
GSC(Homeobox)_FrogEmbryos-GSC-ChIP-Seq(DRA000576)_1D
Hoxd11(Homeobox)_ChickenMSG-Hoxd11.Flag-ChIP-Seq(GSE86088)_1D
Pax7(Paired,Homeobox)_Myoblast-Pax7-ChIP-Seq(GSE25064)_1D
Foxf1(Forkhead)_Lung-Foxf1-ChIP-Seq(GSE77951)_1D
FOXA1(Forkhead)_LNCAP-FOXA1-ChIP-Seq(GSE27824)_1D
FOXA1(Forkhead)_MCF7-FOXA1-ChIP-Seq(GSE26831)_1D
OCT:OCT(POU,Homeobox)_NPC-Brn1-ChIP-Seq(GSE35496)_1D
ASCL1_1D
Hoxc9(Homeobox)_Ainv15-Hoxc9-ChIP-Seq(GSE21812)_1D
Hoxa10(Homeobox)_ChickenMSG-Hoxa10.Flag-ChIP-Seq(GSE86088)_1D
Hoxa9(Homeobox)_ChickenMSG-Hoxa9.Flag-ChIP-Seq(GSE86088)_1D
Hoxd12(Homeobox)_ChickenMSG-Hoxd12.Flag-ChIP-Seq(GSE86088)_1D
Hoxd13(Homeobox)_ChickenMSG-Hoxd13.Flag-ChIP-Seq(GSE86088)_1D
Pdx1(Homeobox)_Islet-Pdx1-ChIP-Seq(SRA008281)_1D
Barx1(Homeobox)_Stomach-Barx1.3xFlag-ChIP-Seq(GSE69483)_1D
Nanog(Homeobox)_mES-Nanog-ChIP-Seq(GSE11724)_1D
Unknown(Homeobox)_Limb-p300-ChIP-Seq_1D
-15 -14 -13 -12 -11 -10 -9 -8 -7 -6 -5 -4 -3 -2 -1 0 1 2 3 4 5 6 7 8 9 10 11 12 13 14 15
4 3 2 1 0
